# TMS-Evoked Responses Are Driven by Recurrent Large-Scale Network Dynamics

**DOI:** 10.1101/2022.06.09.494069

**Authors:** Davide Momi, Zheng Wang, John David Griffiths

**Affiliations:** Krembil Centre for Neuroinformatics, Centre for Addiction & Mental Health, Toronto; Department of Psychiatry & Institute of Medical Sciences, University of Toronto

**Keywords:** recurrence, transcranial magnetic stimulation (TMS), electroencephalography (EEG), connectome, neural mass model, Jansen-Rit, computational model, simulation

## Abstract

A major question in systems and cognitive neuroscience is to what extent neurostimulation responses are driven by recurrent activity. This question finds sharp relief in the case of TMS-EEG evoked potentials (TEPs). TEPs are spatiotemporal waveform patterns with characteristic inflections at ∼50ms, ∼100ms, and ∼150-200ms following a single TMS pulse that disperse from, and later reconverge to, the primary stimulated regions. What parts of the TEP are due to recurrent activity? And what light might this shed on more general principles of brain organization? We studied this using source-localized TMS-EEG analyses and whole-brain connectome-based computational modelling. Results indicated that recurrent network feedback begins to drive TEP responses from ∼100ms post-stimulation, with earlier TEP components being attributable to local reverberatory activity within the stimulated region. Subject-specific estimation of neurophysiological parameters additionally indicated an important role for inhibitory GABAergic neural populations in scaling cortical excitability levels, as reflected in TEP waveform characteristics.

## 1 Introduction

The brain is a complex, nonlinear, multiscale, and intricately interconnected physical system, whose laws of motion and principles of organization have proven challenging to understand with currently available measurement techniques^1^. In such epistemic circumstances, application of systematic perturbations, and measurement of their effects, is a central tool in the scientific armoury^2;3^. For human brains, the technological combination that best supports this non-invasive perturbation-based modus operandi is concurrent transcranial magnetic stimulation (TMS) and electroencephalography (EEG)^4;5^. TMS-EEG allows millisecond-level tracking of stimulation-evoked activity propagation throughout the brain^6;7^, originating from a target region that is perturbed by the secondary electrical currents of a focal (2-2.5cm diameter), brief (∼1ms), and powerful (1.5-2T) magnetic field^8^. Trial-averaged TMS-EEG response waveforms (known as TMS-evoked potentials or TEPs) have been used to elucidate basic neurophysiology in the areas of brain connectivity^9^, axonal conduction delays^10^, and neural plasticity^11^; and also as a clinical biomarker for several pathological conditions such as coma^12^, stroke^13^, depression^14^, obsessive compulsive disorder^15^, and schizophrenia^16^. In addition to this wide variety of basic physiological and clinical applications, TEP measurements speak directly to a central theoretical question at the very heart of systems neuroscience: to what extent does stimulus-evoked neural activity propagate through the brain, via local and/or long-range projections, to affect activity in spatially distant brain regions? In the present paper, we are concerned with this question, and even more so with its equally interesting corollary: to what extent does stimulus-evoked activity propagate *back* from downstream areas to a primary stimulation site. This phenomenon of top-down or cyclic feedback within large-scale brain networks is known as *re-entry* or *recurrence*^17;18;19;20^.

Understanding the contribution of recurrent activity to TEPs, and stimulus-evoked activity in general, is critically important for proper interpretation of TMS-EEG experimental results and design of clinical interventions. In the case of TMS the direct physical and physiological effects at the primary stimulation site of an extracranially-applied magnetic perturbation are reasonably well-understood: secondary electrical currents initially depolarize the membranes of cells in the superficial neural tissue underneath the coil, causing action potentials and an associated local response in the stimulated brain region^21^. Concurrently, this local electrical activation propagates (as some combination of soma-originating and prodromic axon-originating action potentials) along white matter pathways to reach distant cortical and subcortical sites, resulting in predominantly excitatory effects with magnitudes depending on the strength of the anatomical connections^22^. The final EEG-measurable outcomes of this process appear as early (<100ms) and late (>100ms) responses at both the primary stimulation site and a broad set of interconnected brain regions, usually persisting for ∼300ms, and showing reliable characteristic patterns but also high levels of inter-subject variability^23^. A key challenge in interpreting these data is that it is impossible from the TMS-evoked EEG time series alone to disentangle whether TEP waveform components at the primary stimulation sites arise due to a ‘local echo’ - driven only by the recent history of that region, or to a ‘global echo’ - driven by the recurrent activity within the rest of network.

Here we introduce a novel approach to addressing these questions around the physiological basis and spatiotemporal network dynamics of neural activity evoked by noninvasive brain stimulation, using a combination of empirical TMS-EEG data analyses and whole-brain, connectome-based neurophysiological modelling. An overview and conceptual framework is given in Figure 1. The logic proceeds as follows: In a first step, we fit a connectome-based model to individual-subject TEP data, achieving accurate replication of the measured channel- and source-level TMS-EEG patterns. Then, we introduce to the model a series of spatially and temporally specific ‘virtual lesions’ by setting to zero the weights of all connections leaving from and returning to the primary stimulation site, at specific times. These virtual lesions isolate the TMS-stimulated region from the rest of the brain for delineated periods, and allow us to ask what its dynamics would look like with and without recurrent feedback from downstream brain areas. Activity patterns at the stimulated node that are unchanged by a given virtual lesion that suppresses recurrent inputs are thus independent of those inputs, and can be understood as a ‘local echo’ of the stimulation that persists in time for long periods (dozens to hundreds of milliseconds). This framing implies two contrasting potential scenarios for the change in TEP waveform components after introducing a lesion that suppresses recurrent feedback to the stimulation site:

a. TEP components are still observed and entirely or largely unchanged, or
b. TEP components are substantially reduced or disappear

As noted, clear evidence of a) would be consistent with these brain responses being simply a ‘local echo’ of the TMS perturbation, that activates only the stimulated area. In contrast, evidence of b) would imply the local TEP response requires global network reverberation - recurrent activity propagating out from the stimulated region, via its distal interconnected notes, and back again to evoke or to amplify inflections at specific time points post-stimulation.

**Figure 1.**
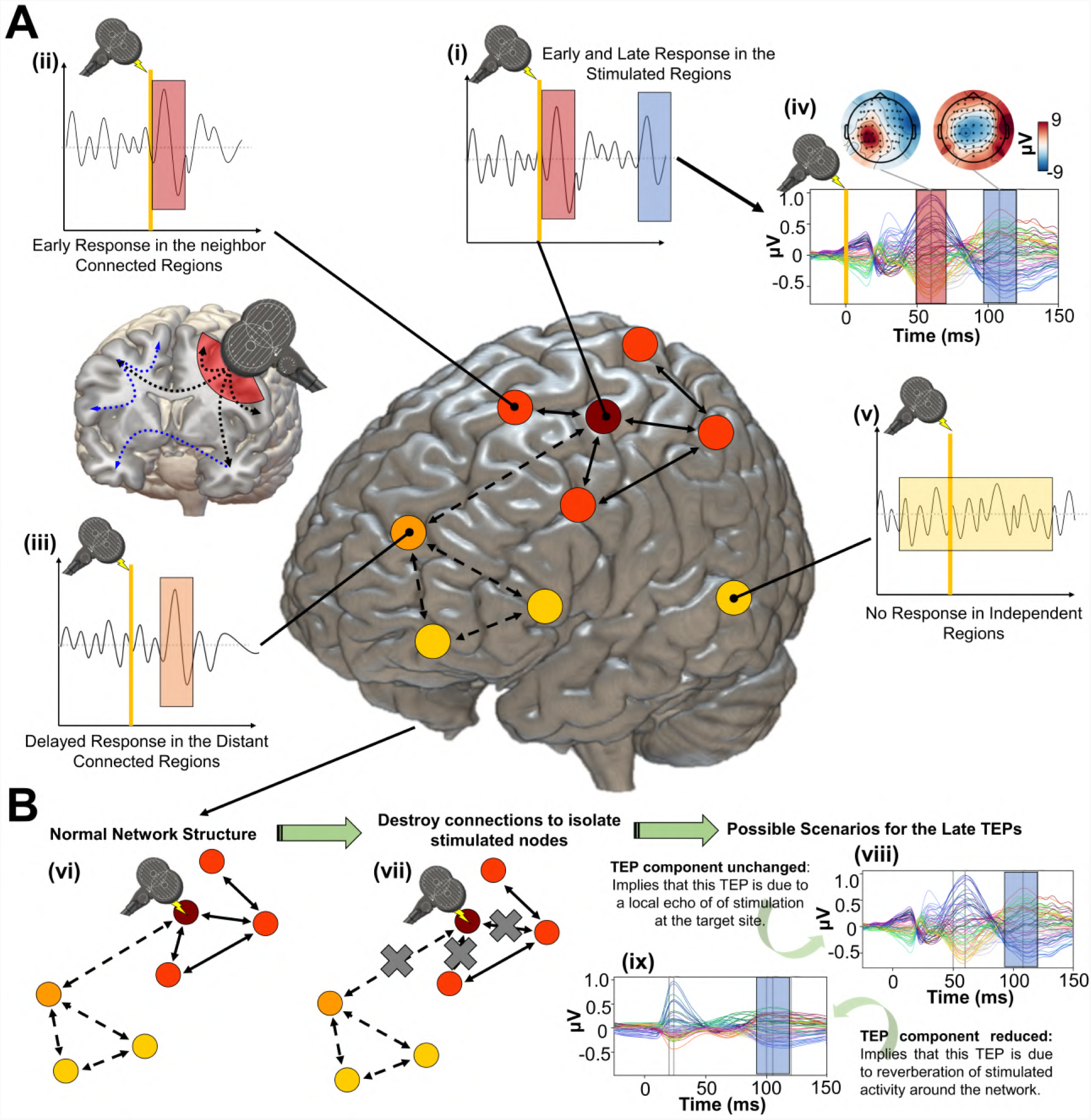
Studying the role of recurrent activity in stimulation-evoked neural responses with computational models. Shown here is a schematic overview of the hypotheses, methodology, and general conceptual framework of the present work. **A)** Single TMS stimulation pulse (*i* - diagram, *iv* - real data) applied to a target region (in this case left M1) generates an early response (TEP waveform component) at EEG channels sensitive to that region and its immediate neighbours (*ii*). This also appears in more distal connected regions such as the frontal lobe (*iii*) after a short delay due to axonal conduction and polysynaptic transmission. Subsequently, second and sometimes third late TEP components are frequently observed at the target site (*i, iv*), but not in non-connected regions (*v*). Our central question is whether these late responses represent evoked oscillatory ‘echoes’ of the initial stimulation that are entirely locally-driven and independent of the rest of the network, or whether they rather reflect a chain of recurrent activations dispersing from and then propagating back to the initial target site via the connectome. **B)** In order to investigate this, precisely timed communication interruptions or ‘virtual lesions’ (*vii*) are introduced into an accurately fitting individual-subject computational model of TMS-EEG stimulation responses (*vi*), and the resulting changes in the propagation pattern (*vii*) are evaluated. No change in the TEP component of interest would support the ‘local echo’ scenario (*viii*), whereas suppressed TEPs would support the ‘recurrent activation’ scenario (*ix*).

For modelling the empirical TMS-EEG TEP data following the investigative line described above, we use a newly-developed numerical simulation approach that draws on recent technical advances from the field of machine learning^24^. Our novel modelling methodology allows accurate and robust individual subject-level TEP waveform fitting, allowing us to present here the first ever subject-specific, cortex-wide, connectome-based neurophysiological model of TEP generation. As we show, this allows us to pose and answer questions around both the shared structure and the well-known inter-subject variability of TMS-EEG responses^25^. We examine the general question of recurrent activity in relation to feedforward and feedback connections to primary stimulation targets, as well as to the broader graph topological structure of the anatomical connectome. Inter-subject variation in estimated physiological parameters offers candidate explanations for TEP phenomena in terms of excitatory/inhibitory population parameters that are consistent with known pharmaco-physiological effects^26^. We argue that this physiologically-based mathematical parameterization of brain stimulation responses offers an important new framework for understanding (and minimizing) inter-subject variability in TMS-EEG recordings for basic scientific and clinical applications.

## 2 Results

### 2.1 Connectome-based neurophysiological models accurately reproduce subject-specific TEP patterns

As an important preliminary to our primary research question, extensive testing confirmed that our new connectome-based neurophysiological model of TMS-EEG responses achieves robust and accurate recovery of TEP waveforms at both the group-average and individual level. This is demonstrated in Figures 2 and 3 for both the EEG channel level and cortical surface source level, respectively. Figure 2A shows empirical and fitted (i.e. simulated, with optimized physiological parameters) TEP waveforms and selected topography maps for three example subjects (for the entire group, see Supplementary Figure S1). It is visually evident in these figures that the model accurately captures several individually-varying features of these time series, such as the timing of the 50ms and 100-120ms TEP components, and the extent to which they are dominated by left/right and temporal/parietal/frontal channels. (For the latter, this can be seen by comparing the line colours in the upper and lower rows of corresponding columns in Figure 2A, and using the channel location references given by the channel colour map on the top left of each TEP plot). Pearson correlations between empirical and simulated TMS-EEG time series confirmed that an excellent goodness-of-fit was observed at the whole-head level (Figure 2B) and individual channel level (Figure 2C), with time-wise permutation tests revealing a significant Pearson correlation coefficient for every electrode. As well as the millisecond-by-millisecond TEP comparisons and the timing of key wwaveform components, we also assessed the accuracy of the model in capturing holistic time series properties. As shown in Figure 2D, a significant positive correlation (*R*^2^ = 80%, *p <* 0.0001) was found between the Perturbational Complexity Index (PCI)^27^ of the simulated and empirical waveforms.

**Figure 2.**
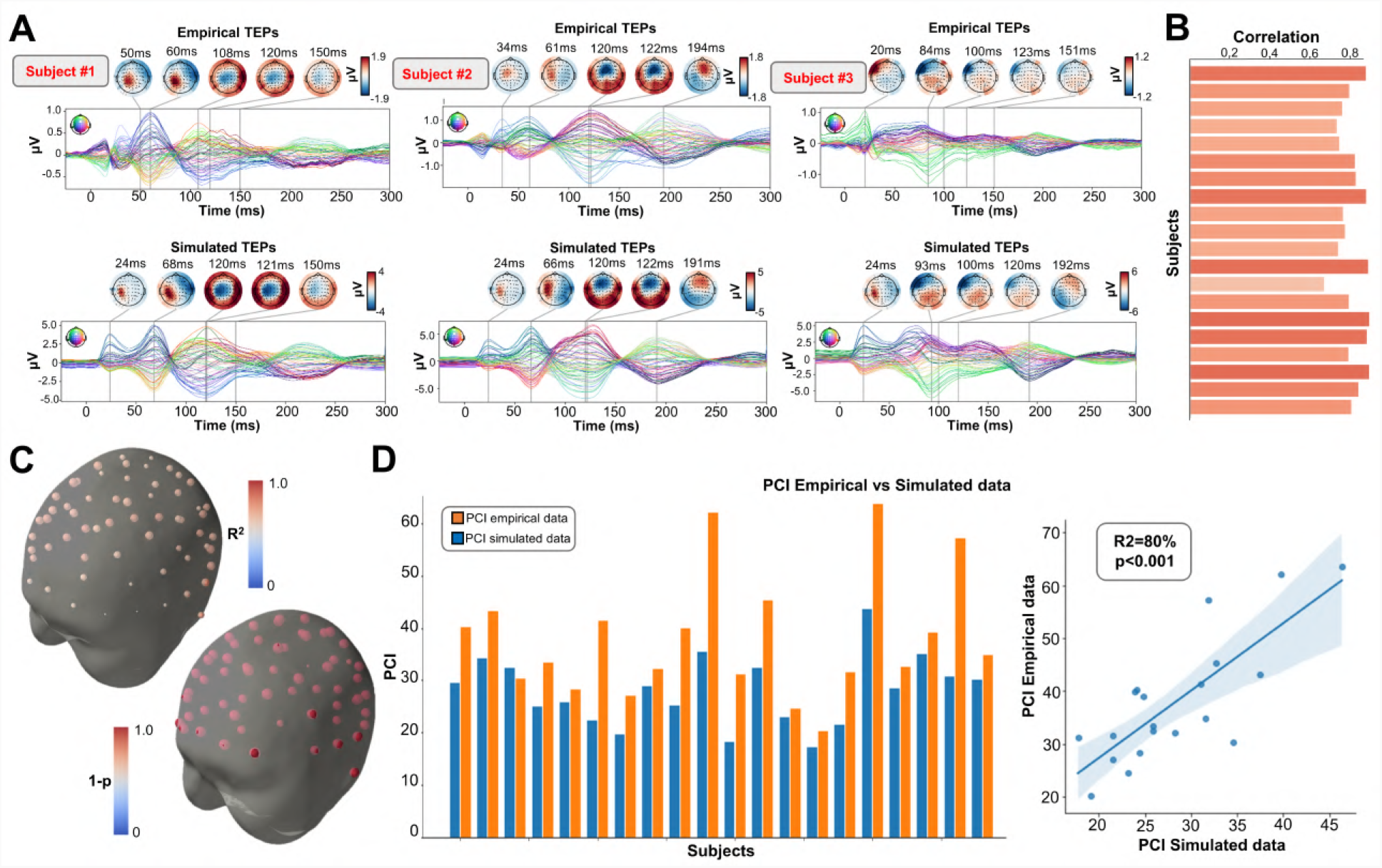
Comparison between simulated and empirical TMS-EEG data in channel space. **A)** Empirical (upper row) and simulated (lower row) TMS-EEG butterfly plots with scalp topographies for 3 representative subjects, showing a robust recovery of individual empirical TEP patterns in model-generated activity EEG time series. **B)** Pearson correlation coefficients between simulated and empirical TMS-EEG time series for each subject. **C)** Time-wise permutation tests result showing the Pearson correlation coefficient (top) and the corresponding significant reversed p-values (bottom) for every electrode. **D)** PCI values extracted from the empirical (orange) and simulated (blue) TMS-EEG time series (left). A significant positive correlation (*R*^2^ = 80%, *p <* 0.0001) was found between the simulated and the empirical PCI (right), demonstrating high correspondence between empirical and simulated data.

**Figure 3.**
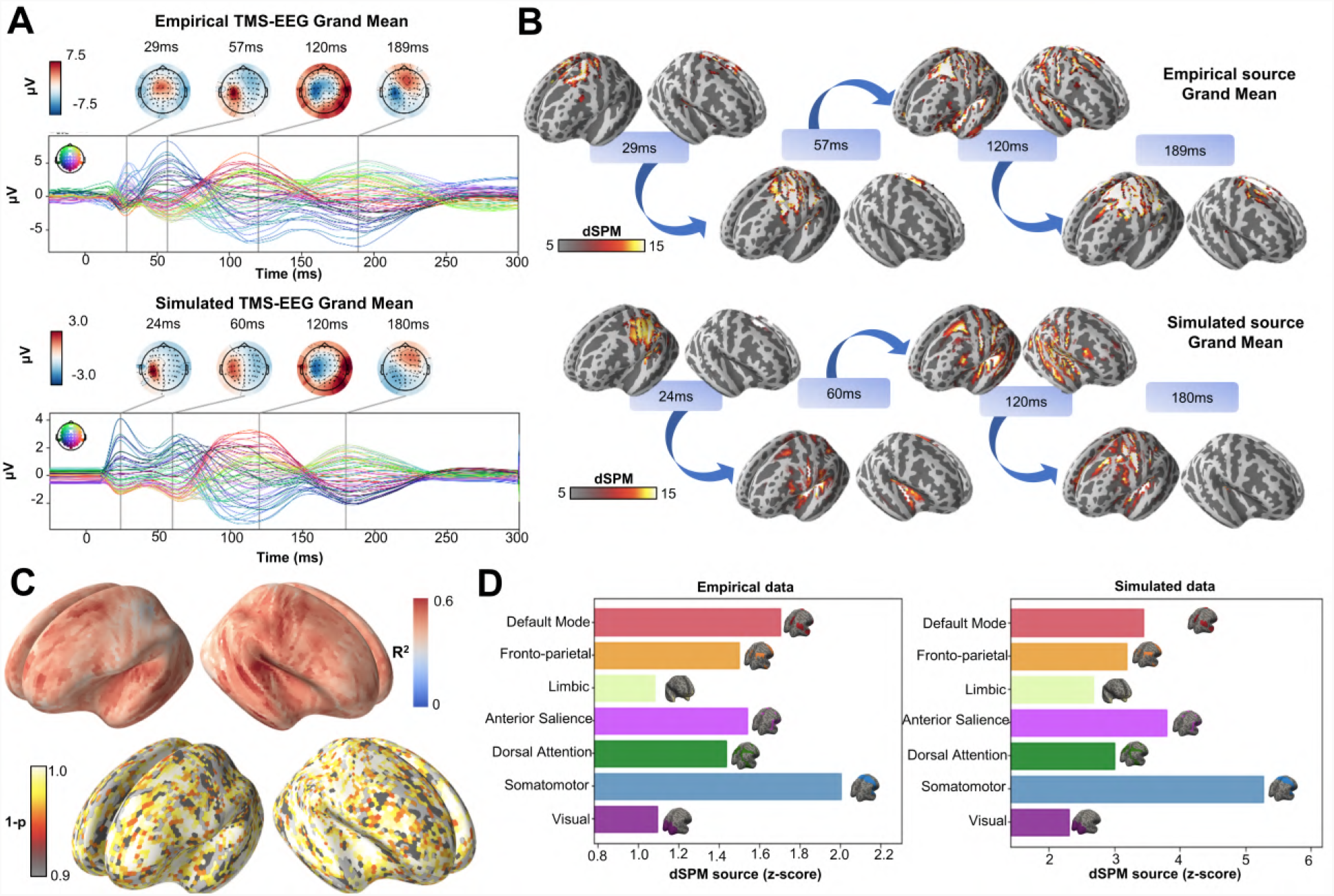
Comparison between simulated and empirical TMS-EEG data in source space. **A)** TMS-EEG time series showing a robust recovery of grand-mean empirical TEP patterns in model-generated EEG time series. **B)** Source reconstructed TMS-evoked propagation pattern dynamics for empirical (top) and simulated (bottom) data. **C)** Time-wise permutation test results showing the significant Pearson correlation coefficient (top) and the corresponding reversed p-values (bottom) for every vertex. **D)** Network-averaged dSPM values extracted for the grand mean empirical (left) and simulated (right) source-reconstructed time series.

Similarly to the single-subject fits, our model also showed accurate recovery of the grand mean empirical TEP waveform when the fitted TEPs were averaged over subjects (Figure 3A). These grand mean channel-level waveforms were further used to assess model fit in brain source space. As can be seen in the topoplots in Figure 3B, the same spatiotemporal activation pattern is observed both for empirical (top row) and model-generated (bottom row) time series. M1 stimulation begins with activation in left motor area at ∼20-30ms, then propagates to temporal, frontal and homologous contralateral brain regions, resulting in a waveform peak at ∼100-120ms. Time-wise permutation tests on these data revealed a significant Pearson correlation coefficient in 75.63% of all vertices (Figure 3C), and again a significant correlation in the simulated and the empirical PCI (*R*^2^ = 80%, *p <* 0.0001).

Finally, significant positive correlations were found between the dynamic Statistical Parametric Mapping (dSPM) values extracted from the 7 canonical Yeo Networks, with stronger correspondences for the primary stimulation target (Somatomotor [SMN], *R*^2^ = 46%, *p* = 0.008) compared to the non-stimulated Networks (Visual [VISN]: *R*^2^ = 38%, *p* = 0.01; Dorsal Attention [DAN]: *R*^2^ = 38%, *p* = 0.004; Anterior Salience [ASN]: *R*^2^ = 38%, *p* = 0.003; Limbic [LIMN]: *R*^2^ = 40%, *p* = 0.01; Fronto-parietal [FPN]: *R*^2^ = 41%, *p* = 0.006; Default Mode [DMN]: *R*^2^ = 43%, *p* = 0.009)). This correspondence between empirical and simulated TEP data in the pattern of their loadings across the canonical networks is clearly visible in the bar charts of Figure 3D.

### 2.2 Later TEP responses are driven by recurrent large-scale network dynamics

Having established the accuracy of our model at replicating TEP waveforms across a wide range of subject-specific shapes (Figures S1, 2, 3), we now turn to the central question of the present study, as laid out in Figure 1B. Shown in Figure 4 are effects on the simulated TEP of virtual lesions to recurrent incoming connections of the main activated regions at 20ms, 50ms, and 100ms after single-pulse TMS stimulation of left M1. The leftmost column of Figure 4, which shows the simulated data grand average with no virtual lesion (re-plotting the data from the second row of Figure 3B), serves as a reference point for other three columns. A key result here is that there is a reduction of source activation at the 50-100ms time window in the central two panels (lesions at 20ms and 50ms, respectively), as compared to the rightmost (lesion at 100ms) and leftmost (no lesion) panels. This reduction demonstrates the critical importance of network recurrence in generating later TEP components. These effects were evaluated statistically by extracting dSPM loadings from the source activity maps for each of the 7 canonical Yeo networks^28^ and entering them into an ANOVA with factors of NETWORK and TIME OF DAMAGE. Significant main effects were found for both NETWORK (F_(6,114)_ = 114.73, p < 0.0001, 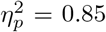) and TIME OF DAMAGE (F_(3,57)_ = 254.05, p < 0.0001, 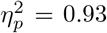) - indicating that the effects of virtual lesions vary depending on both the time administered and the site administered to, as well as the combination of these factors (significant interaction NETWORK*TIME OF DAMAGE (F_(18,342)_ = 23.79, p < 0.0001, 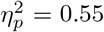). The specific results described above, with significant TEP suppression at the stimulation site occurring in the early and late TEP components for specific time windows, was verified through extensive post-hoc t-tests (see Supplementary Results Section 2.1).

**Figure 4.**
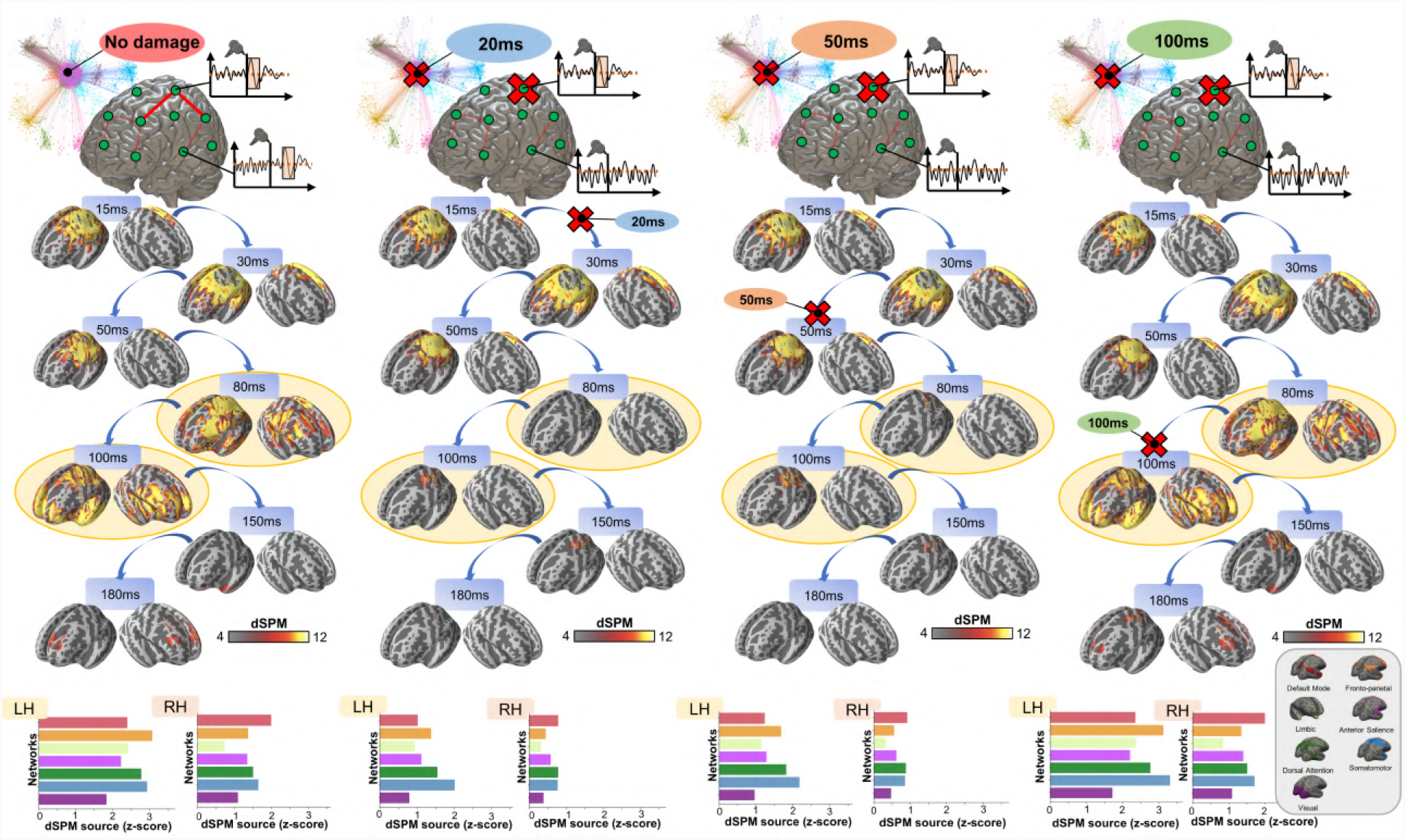
Removing recurrent connections from stimulated target nodes suppresses their late TEP activity. We found that TMS-evoked propagation dynamics in the model change significantly depending on the specific time that a virtual lesion is applied (highlighted orange circle). Specifically, early significant reductions in the TMS-evoked activity (50ms-100ms time window) were found when important connections were removed at 20ms (blue) and 50ms (orange) after the TMS pulse, as compared to both a later virtual lesion (100ms green) and no damage (red) conditions. This results is demonstrated also for network-based dSPM values (bottom row) extracted for all four conditions.

### 2.3 TMS-evoked activity propagation strongly depends on highly connected brain regions

After demonstrating the importance for TEPs of recurrent feedback into the primary stimulation regions, we next asked whether the activity propagation patterns observed in TMS-EEG also depend on more global topological features of the anatomical connectome. In order to assess this, we performed the same time-windowed virtual lesion analyses for two ATTACK TYPE conditions: targeted attack where the most important nodes in the brain network’s graph structure were damaged; random attack where 1000 simulations were run and the nodes selected for removal were randomly chosen. As shown in Figure 5B, by analyzing the PCI values at the channel level, significant main effects of ATTACK TYPE (F_(1,19)_ = 62.01, p < 0.0001, 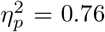) and TIME OF DAMAGE (F_(2,38)_ = 23.76, p < 0.0001, 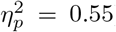) were observed, as well as a significant interaction ATTACK TYPE*TIME OF DAMAGE (F_(2,38)_ = 22.63, p < 0.0001, 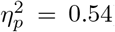). This indicates that both the time and the type of the virtual lesion highly affect the propagation of the activity elicited by a short TMS stimulation. Specifically, considering the type of the lesion, targeted attack conditions significantly reduced EEG time series complexity compared to random attack conditions. Conversely, considering the time of the lesion, the effects of early targeted attacks (20ms and 50ms) are significantly higher compared to later lesion times (100ms).

**Figure 5.**
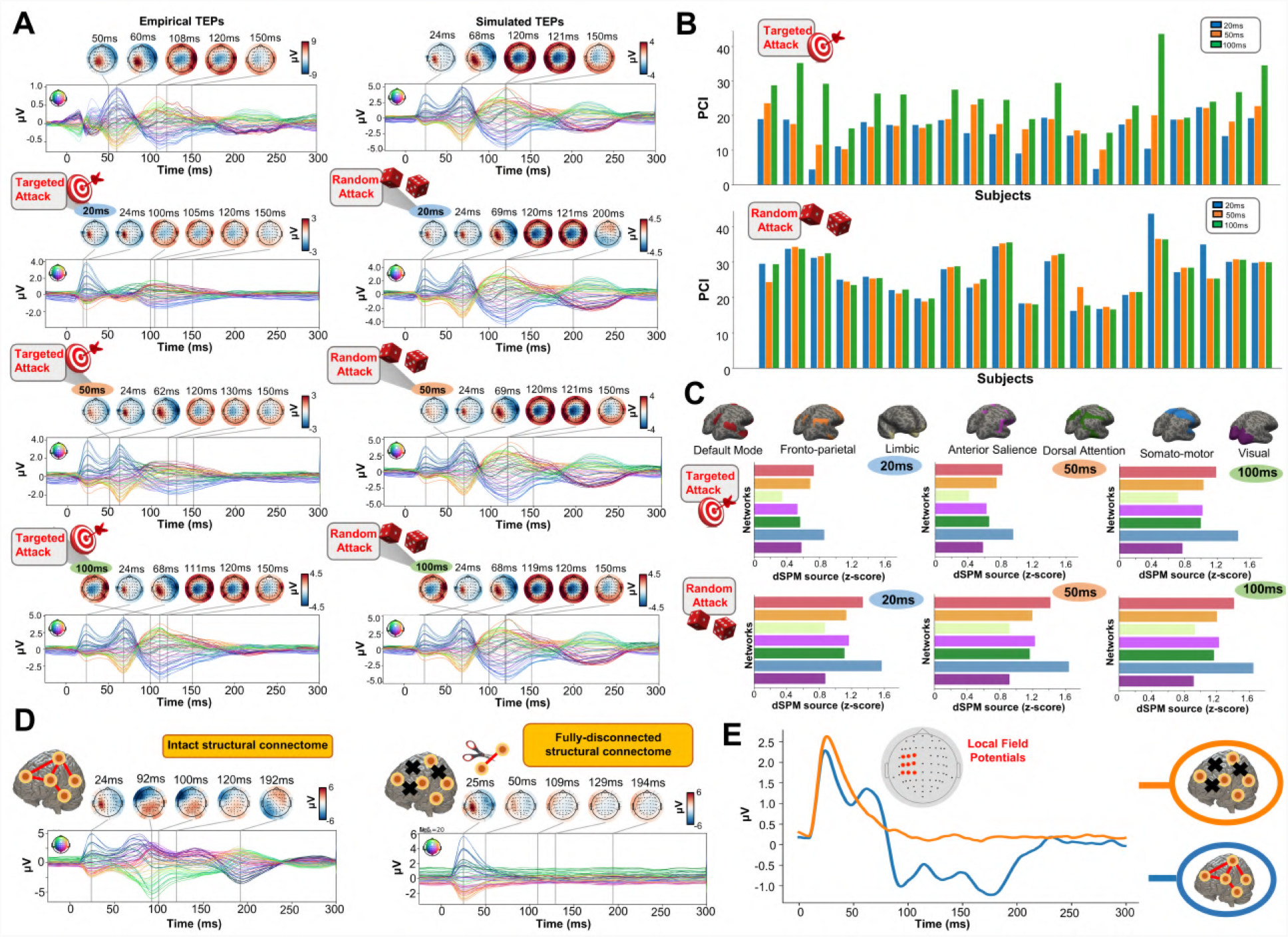
TMS-evoked activity propagation depends on connectome hubs. **A)** Effect of two anatomical connectivity-based lesion strategies (random vs targeted) and time of damage (20ms: blue; 50ms: orange; 100ms: green) on TMS-EEG dynamics for one representative subject. Overall, targeted attack (left column) significantly compromised the propagation of the TMS-evoked signal compared to the random attack (right column) condition. Moreover, the EEG dynamics were significantly affected by early (20ms: blue and 50ms: orange) compared to late (100ms: green) virtual lesions. **B)** PCI scores extracted for targeted (top) and random (bottom) attack and for the different time of damage conditions. A gradient in the PCI scores was found for the targeted attack condition, where earlier lesions were associated with the lower complexity and later ones with higher complexity. Conversely, in the random attack condition, PCI did not differ significantly for different lesion times. **C)** Grand average dSPM values extracted from source reconstructed TMS-EEG surrogate data for targeted (top) and random (bottom) attack and for the three different time of damage conditions. Similarly to the channel-level results, source activity was significantly reduced for targeted attack compared to random lesions. A significant decrease in the source-reconstructed activity was found after early compared to late connectome damage. Interestingly, these effects were highest in the network receiving the initial stimulation, namely the sensorimotor network. **D)** Demonstration of the network recurrence-based theory for one representative subject. Simulation of TMS-EEG dynamics run using the intact (left) and fully-disconnected (right) anatomical connectome. In the latter case the external perturbation generates a local response which reverberates locally and terminates after ∼50ms. This demonstrates again how later TEPs are driven by recurrent network dynamics. **E)** Local Mean Field Power (LMFP) at the stimulation site for intact (blue line) and fully-disconnected (orange line) anatomical connectome.

To gain further insight into the anatomy of these effects, we then evaluated the effects of different virtual lesions (type and timing) on the source reconstructed signal for each of the 7 Yeo et al. functional networks^28^. Network dSPM values at source level (Figure 5C) showed significant main effects of NETWORK (F_(6,114)_ = 42.99, p < 0.0001, 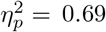) ATTACK TYPE (F_(1,19)_ = 46.91, p < 0.0001, 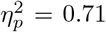), and TIME OF DAMAGE (F_(2,38)_ = 44.55, p < 0.0001, 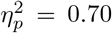), as well as a significant double interaction ATTACK TYPE*TIME OF DAMAGE (F_(2,38)_ = 27.12, p <0.0001, 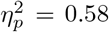), NETWORK*TIME OF DAMAGE (F_(12,228)_ = 10.62p < 0.0001, 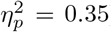), and a significant triple interaction NETWORK*ATTACK TYPE*TIME OF DAMAGE (F_(12,228)_ = 6.28, p < 0.0001, 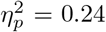). The significant main and double effects here again show that different networks are affected at different times and to different magnitudes by virtual lesion connectivity disruptions, underscoring the pivotal role of both time and space for shaping the propagation of the brain activity induced by an external perturbation. For further details on the post-hoc analyses pertaining to these ANOVA results please refer to Supplementary Results Section 2.2. For a representation of the nodes removed, please refer to Supplementary Figures S2. For a comprehensive overview of individual changes in TMS-EEG dynamics after different lesions, please refer to Supplementary Figures S3, S4 and S5.

### 2.4 Inhibitory synaptic activity accounts for inter-subject differences in TEP waveforms

One of the key advantages of physiologically-based brain modelling is the potential for making meaningful associations between major empirical data features and the physiological constructs instantiated in the model’s parameters. We explored this by examining the relationship between TEP waveform components and physiological parameters of the Jansen-Rit model. To do this, singular value decompositions (SVDs) were performed on the channel x time TEP waveform matrices for both empirical and simulated data. The left and right singular vectors from this decomposition respectively define the temporal and spatial expression of the channel-level TEP *eigenmodes*. The spatial part of each eigenmode takes the form of a loading pattern over channels that can be represented as a topoplot. As with the TEP waveform and PCI comparisons, this procedure also yielded high spatial similarity between empirical and simulated grand average data (Figure 6A), as well as similar levels of variance explained (74.14% and 66.96% cumulatively by the first two right SVD eigenvectors in simulated and empirical data, respectively). Inspecting the temporal peaks in the left singular vectors for the first two eigenmodes revealed that the first was maximally expressed in empirical [/simulated] data at 72ms[/70ms], and the second at 115ms [/117ms] post-stimulus. Thus the first two eigenvectors of TEP waveform correspond quite closely to the canonical ∼50ms and ∼100ms TEP waveform components. As shown in Figure 6B, a significant negative correlation was found between the synaptic time constant of the Jansen-Rit inhibitory population and the amplitude of the first eigenmode at its peak (*R*^2^ = 27%, *p* = 0.02). Interestingly, we also observed a significant positive correlation between this parameter and the second eigenmode at its peak (*R*^2^ = 28%, *p* = 0.02). For a comprehensive overview of individual timing and topographies of the first two eigenmodes, please refer to Supplementary Figures S6.

**Figure 6.**
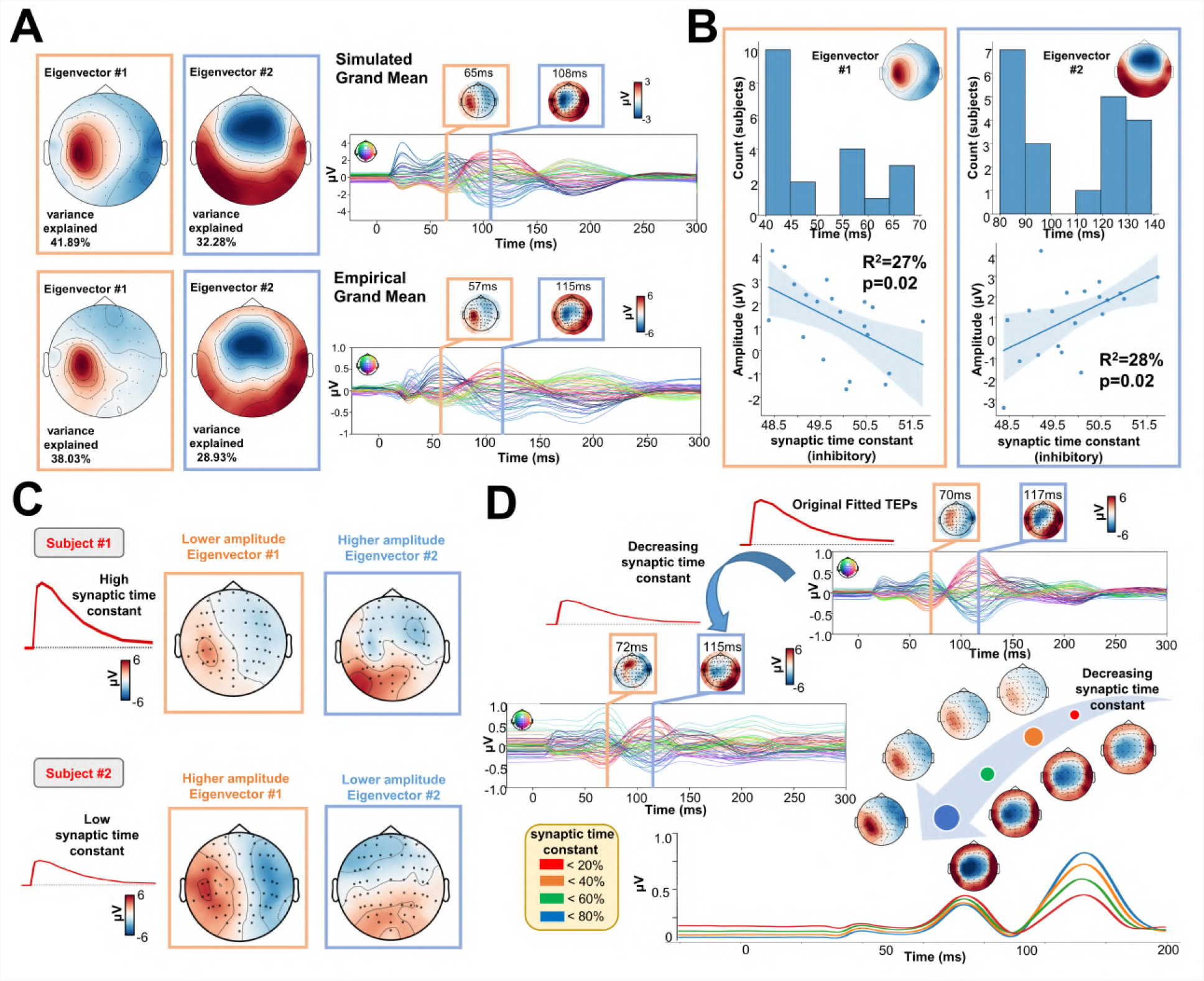
Synaptic time constants of inhibitory neural populations affect early and late TEP amplitudes. **A)** Singular value decomposition (SVD) topoplots for simulated (top) and empirical (bottom) TMS-EEG data. Results revealed that the first (orange outline) and the second (blue outline) SVD eigenmodes were located ∼65ms and ∼110ms after the TMS pulse, respectively. **B)** First and second SVD temporal eigenmode latencies and amplitudes were extracted for every subject, and the distribution plots (top row) show the time window where highest cosine similarity with the SVD spatial eigenvectors was found. Scatter plots (bottom row) show a significant negative (left) and positive (right) correlation between the synaptic time constant of the inhibitory population and the amplitude of the the first and second eigenvector. **C)** Model-generated first and second SVD eigenmodes for 2 representative subjects with high (top) and low (bottom) estimated values for the synaptic time constant of the inhibitory population. The topoplots show that the magnitude of the synaptic time constant is closely coupled to the the amplitude of the individual first and second SVD modes. **D)** Model-generated TMS-EEG data were run using the optimal (top right) or 85% decreased (central left) value for the synaptic time constant of the inhibitory population. The bottom right panel shows absolute values for different magnitudes of this parameter. Results show an increase in the amplitude of the first, early, and local TEP component; and a decrease of the second, late, and global TEP component, as a function of the inhibitory synaptic time constant.

## 3 Discussion

Using our novel computational framework for personalized TMS-EEG modelling, in this work we have presented new insights into the role of recurrent activity in stimulation-evoked brain responses. Characterizing these phenomena at a mechanistic level is important not only as a basic question in systems and cognitive neuroscience, but also as a foundation for clinical applications concerned with changes in excitability and connectivity due to neuropathologies or interventions.

We employed a ‘virtual dissection’ approach^29^ to study the extent to which model-generated TMS-evoked stimulation patterns at the primary stimulation site relied on recurrent incoming connections from the rest of the brain, and at what times. These *in-silico* interventions resulted in substantial reductions in TMS-evoked activity when pivotal connections were inactivated. Specifically, compared to late (100ms after the TMS pulse) virtual lesions, and compared to the control condition where no damage was applied, early (20ms, 50ms) damage of essential nodes’ afferent and efferent pathways significantly reduced the amplitude of the 100ms TEP component at the stimulation site (left M1) and its neighbouring regions (Figure 4). In these early lesion conditions some residual activity in the left M1 area was still observed at around 100ms, indicating that a local echo of the TMS stimulus does indeed persist for tens to hundreds of milliseconds after stimulation. However, this purely locally-driven activity was low in amplitude, and does not appear to be the principal source of the commonly studied 100ms TEP components in TMS-EEG recordings. In additional to recurrence at the stimulation site, we can also see that amplification of the TMS-evoked stimulation response occurs via network spreading and recruitment. Early lesions also compromised the propagation of the TMS-evoked activity to the contralateral homologue of the stimulated region (i.e. right M1), as well as bilateral frontal and parietal regions. This result clarifies not only *that* TMS-evoked activity in those regions depends on the presence of those specific cross-hemispheric and parieto-frontal pathways in the network, but also *when* propagation along them is critical for the subsequent response. Finally, in contrast to the 100ms TEP component, the 50ms TEP component at the target site was largely unaffected by lesions to recurrent connections at 20ms and 50ms, indicating that this earlier part of the the canonical TMS-EEG response can be attributed solely to the local impulse-response characteristics of a patch of cortical tissue.

Our results, and the framework for investigating such questions that we are introducing here, have clear and practical relevance to basic and clinical TMS-EEG research, but also have broader implications for the scientific understanding of functional brain organization. Variations on the concept of recurrence in systems neuroscience go back many decades, and have been developed in a wide number of areas and with a wide number of labels, including ‘re-entry’, ‘reverberation’, ‘feedback’, ‘top-down control’, ‘predictive coding’, ‘functional/effective connectivity’, etc^18;19;30;12;3;17^. These framings vary a great deal on dimensions such as the spatial/temporal scale, role of corticothalamic interactions, association with cognitive functions, association with global brain state, level of physiological detail / abstraction, etc. In all these cases however the central shared intuition is that information or activity flows between network elements in the brain are bidirectional, but that the primary direction of travel may fluctuate dynamically over time. For example, the response of the visual system to images - a sensory stimulation-evoked response that is similar in many ways to electromagnetic stimulation-evoked responses - is widely understood to involve a period of feedforward activity propagation hierarchically up the ventral visual stream, followed rapidly by recurrent top-down feedback^31;32;33^. Moreover, in vitro recordings have shown how magnetic pulse delivered to a single ganglion cell generates a local early response that terminates after few ms^34^ depicting a scenario similar to Figure 5D-E. The connectome-based neurophysiological modelling approach presented here could easily be deployed to investigate similar questions in these and other areas such as visual cognitive neuroscience or consciousness research, where feedback and recurrence play a central explanatory role in current theories.

The mathematical and theoretical neuroscientific context that has particularly informed the present study owes much to the ideas of Walter Freeman^3^ of Andreas Spiegler and colleagues^30^. Freeman’s hierarchical ‘K Set’ framework^3^ offers a rigorous technical and qualitative analysis of neuronal dynamics in systems progressing in complexity from a single excitatory neural population (K0e set), to ones with self-, uni-directional, and bi-directional excitation an inhibition (KI sets), and eventually adding network-level interactions and feedback (KII and KIII sets). Notably, Freeman’s analysis provides both physiological and mathematical motivation for the central premise of our argument - that a local patch of cortical neural tissue can generate TEP-like damped oscillatory responses to a brief stimulation, without the need for feedback from other brain regions. (This is also an implicit premise in all studies using second-order differential equations to model sensory-evoked potentials, such as Freeman^3^, Jansen-Rit^35^, David^36^, and ourselves here.) In these terms then, the questions we have posed and addressed are whether the 50ms and 100ms TEP components at the stimulation site represent KI set or KII set ensemble behaviour. Complementing this, the nature of recurrent activity at the level of whole-brain connectome networks in particular is expressed more sharply in the work of Spiegler et al.^30^, who emphasizes how feedback loops within the connectome can lead to re-entrant activity, the result of which is to produce longer-lasting and temporally more complex evoked responses - consistent with our findings here. These authors also discovered from an exhaustive investigation comprising 37,000 simulation runs over 190 different stimulation targets that persistent, long-lasting activations tend to propagate within canonical resting-state networks. Interestingly, this prediction was later confirmed in our own experimental TMS-EEG work^37;25^, which demonstrated that the TEPs mainly propagate within distal cortical regions belonging to the same network. For example, stimulation of parietal default-mode network (DMN) nodes resulted in widespread sustained activity across the parietal, temporal, and frontal lobes - but this activity was primarily to be found within other DMN regions. The same result was also observed for nearby stimulation of dorsal attention network (DAN) nodes. More recently we obtained a similar result with anatomical connectivity^9^, namely that network-level anatomical connectivity is more relevant than local and global brain properties in shaping TMS signal propagation after the stimulation of two resting-state networks (again DMN and DAN). Whilst we did not study DMN or DAN stimulation in the present study, it can be seen from the Yeo network loadings in Figures 3-5 that our results are also consistent with these experimental observations, with the somatomotor network dominating for all our simulated M1 stimulations. Extending the present results to TEP measurements from additional target sites both anterior and posterior to the M1 target studied here is an important priority for future work with this model.

In addition to our scientific conclusions on the nature of recurrent activity in stimulation-evoked brain dynamics, the present work offers several technical advances over previous contributions in a number of areas. Our model is to our knowledge the first connectome-based neurophysiological model for TMS-EEG that demonstrates accurate single-subject reconstruction of TEP waveforms at the sensor and source level. Related work has focused on stimulation-evoked functional connectivity patterns^30^ and stimulation-evoked time-frequency responses^38^ within either large or small-scale networks. Most notably and recently, Bensaid and colleagues^39^ proposed a whole-brain model of TMS-EEG TEP waveforms, with a focus on the sleep/wake differences in TMS-EEG responses studied by Casali^27^, Massimini^12^, and others. Bensaid et al’s model includes extensive ‘horizontal’ corticothalamic connectivity, which we elected not to replicate in the present model for reasons of tractability, but may add in future iterations. None of the above studies, or indeed any published work to date to our knowledge, achieve the level of accuracy for single subject TEP waveform fits that we show here. Our model’s success on this front is owed in large part to our decision to formulate and implement the Jansen-Rit connectome network differential equations in the widely-used machine learning library PyTorch^40^. We have recently discussed and demonstrated the advantages of deep learning-based computational architectures for neurophysiological model simulation and parameter estimation^24^. In the present study this precision was critical for addressing our research questions, which centred on the timing and amplitudes of well-defined TEP waveform components. These components can be found in most or all subjects, but vary considerably in their shapes and exact timings.

One example of the utility of this new model-fitting framework can be seen in our results in Figure 6, where we identified trends over subjects in the relationship of estimated model parameters to individual variation in TEP waveform features. Through these analyses we found, in an entirely data-driven fashion, that the synaptic time constant of the inhibitory Jansen-Rit population is a strong predictor of the amplitude of early (P60) and late (N100) TEP components. This is consistent with the finding of increased TEP amplitudes following application of paired-pulse TMS protocols known to effect inhibition (or reduced excitability)^41^. Similarly, pharmacological intervention studies have shown that GABA_B_ receptor agonists (benzodiazepine) decrease N100 component amplitude, suggesting that this component is driven by GABA_B_ receptor-mediated inhibition. Whilst further research will be needed to explore and verify this hypothesis, its generation via the combination of data-driven model fitting and theoretically-informed brain network simulations offers a promising new approach for interpretation of TMS-EEG experiments, and neurophysiological research more broadly.

## 4 Material and Methods

### 4.1 Overview of approach

The analyses conducted in the present study consist of four main components: i) TMS-EEG evoked response source reconstructions, ii) construction of anatomical connectivity priors for our computational model using diffusion-weighted MRI (DW-MRI) tractography, iii) simulation of whole-brain dynamics and stimulation-evoked responses with a connectome-based neural mass model, and iv) fitting of the model to individual-subject TMS-EEG data. A schematic overview of the overall approach given in Figure 7.

**Figure 7.**
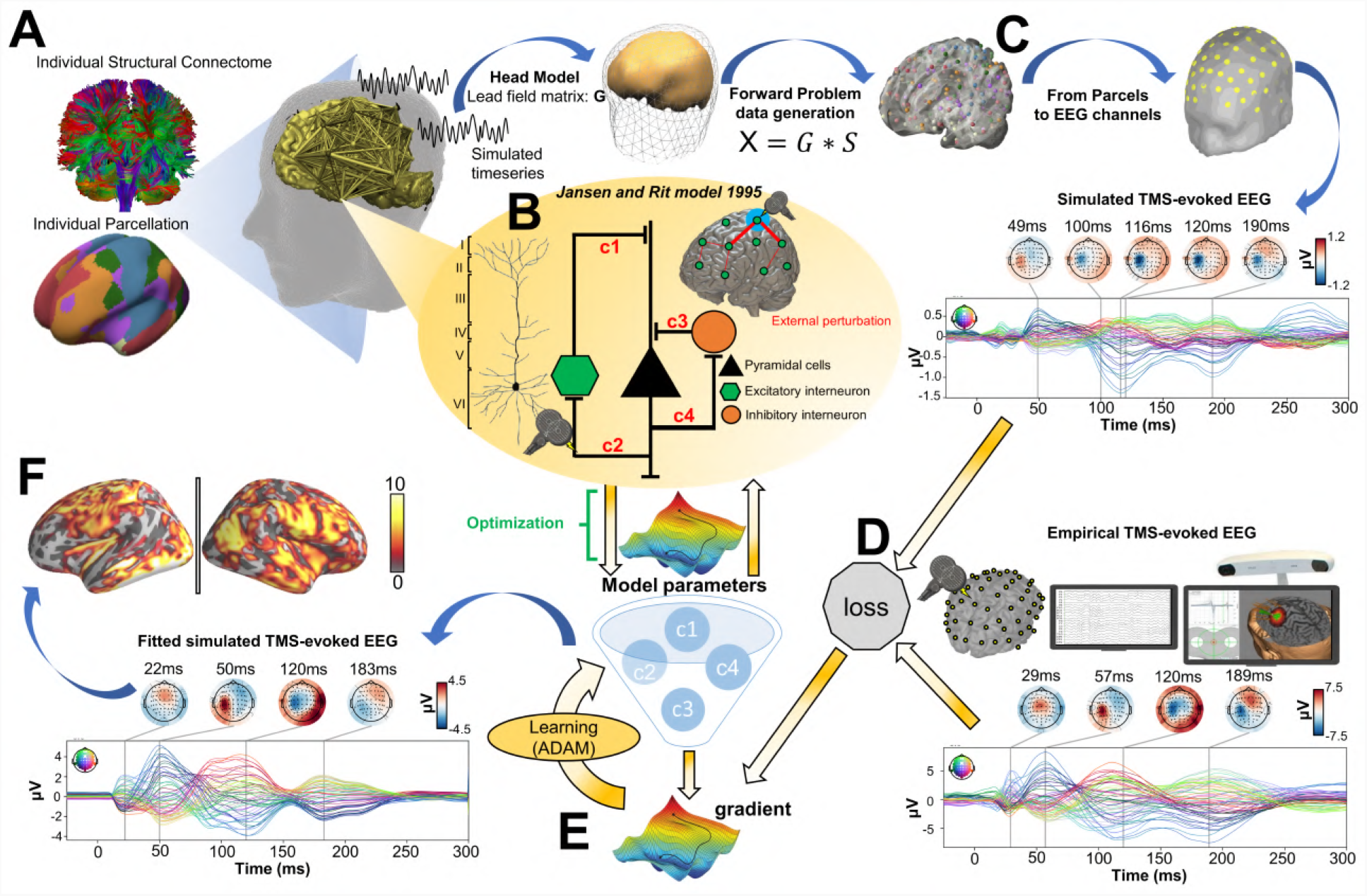
Methodological workflow for subject-specific connectome-based neurophysiological modelling of TMS-EEG TEPs. **A)** DW-MRI tractography was computed from a sample of healthy young individuals from the Human Connectome Project (HCP) Dataset^42^, and then averaged to give a grand-mean anatomical connectome. The 200-parcel Schaefer atlas^43^ was used, which usefully aggregates its 200 brain regions into 7 canonical functional networks (Visual network: VISN, Somatomotor network: SMN, Dorsal attention network: DAN, Anterior salience network: ASN, Limbic network: LIMN, Fronto-parietal network: FPN, Default mode network: DMN). These parcels were mapped to the individual’s FreeSurfer parcellation using spherical registration^44^. Once this brain parcellation covering cortical structures was extrapolated, it was then used to extract individual anatomical connectomes. **B)** The Jansen-Rit model^35^, a neural mass model comprising pyramidal, excitatory interneuron, and inhibitory interneuron populations was embedded in every parcel for simulating and fitting neural activity time series. The TMS-induced depolarization of the resting membrane potential was modelled by a perturbing voltage offset to the mean membrane potential of the excitatory interneuron population. **C)** A lead-field matrix was then used for moving the parcels’ time series into channel space and generating simulated EEG measurements. **D)** The goodness-of-fit (loss) was calculated as the cosine similarity between simulated and empirical TMS-EEG time series. **E)** Utilizing the autodiff-computed gradient^45^ between the objective function and model parameters, model parameters were optimized using the ADAM algorithm^46^. **F)** Finally, the optimized model parameters were used to generate the fitted, simulated TMS-EEG activity, for which we report comparisons with the empirical data at both the channel and source level using conventional statistical techniques.

### 4.2 TMS-EEG data and source reconstruction

The TMS-EEG data used in this study were taken from an open dataset collected and provided to the community by the Rogasch group (figshare.com/articles/dataset/TEPs-_SEPs/7440713), where high-density EEG was recorded following a stimulation of primary motor cortex (M1) in 20 healthy young individuals (24.50±4.86 years; 14 females), and in which state-of-the-art preprocessing had already been applied. For details regarding the data acquisition and the preprocessing steps please refer to the original paper of Bibiani et al.^47^. All TMS-evoked EEG source reconstruction was performed using the MNE software library^48^ (mne.tools/stable/index.html) running in Python 3.6. First, the watershed algorithm was used to generate the inner skull, the outer skull and the outer skin surface triangulations for the ‘fsaverage’ template. Then the EEG forward solution was calculated using a three compartment boundary-element model^49^. Noise covariance was estimated from individual trials using the pre-TMS (from -1000ms to -100ms) time window as baseline. The inverse model solution of the cortical sources was performed using the dSPM method with current density^50^ and constraining source dipoles to the cortical surface. The resulting output of EEG source reconstruction was the dSPM current density time series for each cortical surface location.

### 4.3 Neuroimaging data and definition of connectome weight priors

The whole-brain model we fit to each of the 20 subjects’ TMS-EEG consists of 200 brain regions, connected by weights of the anatomical connectome. We set strong priors on the connection weights, such that individual fits allow for small adjustment of these values. To obtain population-representative values for these connectivity priors, we ran diffusion-weighted MRI tractography reconstructions across a large number of healthy young subjects and averaged the results. For these analyses we used structural neuroimaging data of 400 healthy young individuals (170 males; age range 21-35 years), taken from the Human Connectome Project (HCP) Dataset (`humanconnectome. org/study/hcp-young-adult`)^42^. DW-MRI preprocessing was run in Ubuntu 18.04 LTS, using tools from the FMRIB Software Library (FSL 5.0.3; www.fmrib.ox.ac.uk/fsl)^51^, MRtrix3 (www.MRtrix.readthedocs.io)^52^ and FreeSurfer 6.0^53^. All images used were already corrected for motion via FSL’s EDDY^54^ as part of the HCP minimally-preprocessed diffusion pipeline^55^. The multi-shell multi-tissue response function^56^ was estimated using constrained spherical deconvolution^57^. T1-weighted (T1w) images, which were already coregistered to the b0 volume, were segmented using the FAST algorithm^58^. Anatomically constrained tractography was employed to generate the initial tractogram with 10 million streamlines using second-order integration over fiber orientation distributions^59^. Then, the spherical-deconvolution informed filtering of tractograms (SIFT2) methodology was applied^60^, in order to provide more biologically accurate measures of fibre connectivity. Brain regions or network nodes were defined using the 200-region atlas of Schaefer et al.^43^, which was mapped to each individual’s FreeSurfer surfaces using spherical registration^44^. This atlas additionally provides categorical assignments of regions into 7 canonical functional brain networks (Visual network: VISN, Somatomotor network: SMN, Dorsal attention network: DAN, Anterior salience network: ASN, Limbic network: LIMN, Fronto-parietal network: FPN, Default mode network: DMN). Using this atlas in combination with the filtered streamlines, 200×200 two anatomical connectivity matrices were extracted, with matrix elements representing the number of streamlines and the fiber lenght connecting each pair of regions, respectively. These connectomes for the 400 HCP subjects were then averaged, yielding a healthy subject population-representative connectome matrix. Finally, this matrix was prepared numerically for physiological network modelling by rescaling values by first taking the matrix Laplacian, and second by scalar division of all entries by the matrix norm.

### 4.4 Large-scale connectome-based neurophysiological brain network model

As previously described, a brain network model comprising 200 cortical areas was used to model TMS-evoked activity patterns, where each network node represents population-averaged activity of a single brain region according to the rationale of mean field theory^61^. We used the Jansen-Rit (JR) equations to describe activity at each node, which is one of the most widely used neurophysiological models for both stimulus-evoked and resting-state EEG activity measurements^35;36;62^. JR is a relatively coarse-grained neural mass model of the cortical microcircuit, composed of three interconnected neural populations: pyramidal projection neurons, excitatory interneurons, and inhibitory interneurons. The excitatory and the inhibitory populations both receive input from and feed back to the pyramidal population but not to each other, and so the overall circuit motif (Figure 7B) contains one positive and one negative feedback loop. For each of the three neural populations, the post-synaptic somatic and dendritic membrane response to an incoming pulse of action potentials is described by the second-order differential equation

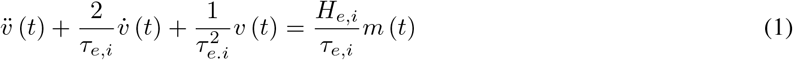

which is equivalent to a convolution of incoming activity with a synaptic impulse response function

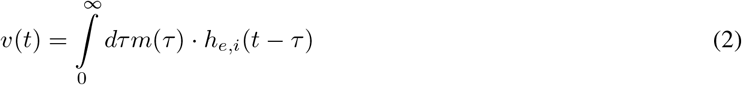

whose kernel *h*_*e,i*_(*t*) is given by

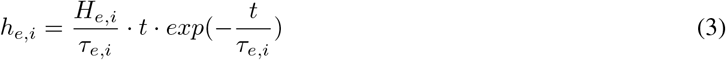

where *m*(*t*) is the (population-average) presynaptic input, *v*(*t*) is the postsynaptic membrane potential, *H*_*e,i*_ is the maximum postsynaptic potential and *τ*_*e,i*_ a lumped representation of delays occurring during the synaptic transmission.

This synaptic response function, also known as a pulse-to-wave operator^3^, determines the excitability of the population, as parameterized by the time constants *τ*_*e*_ and *τ*_*i*_, which are of particular interest in the present study. Complementing the pulse-to-wave operator for the synaptic response, each neural population also has wave-to-pulse operator^3^ that determines the its output - the (population-average) firing rate - which is an instantaneous function of the somatic membrane potential that takes the sigmoidal form

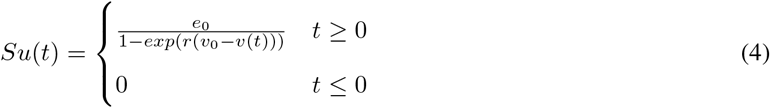

where *e*_0_ is the maximum firing rate, *r* is the steepness of the sigmoid function, and *v*_0_ is the postsynaptic potential for which half of the maximum firing rate is achieved.

In practice, we re-write the three sets of second-order differential equations that follow the form in Equation 1 (one for each population in the JR circuit) as three interconnected pairs of coupled first-order differential equations, and so the full JR system for each individual cortical area *j* ∈ *i* : *N* in our network of *N* =200 regions is given by the following six equations:

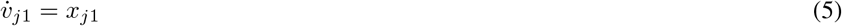

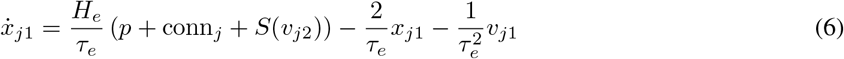

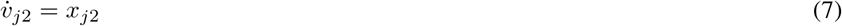

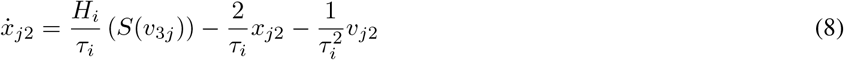

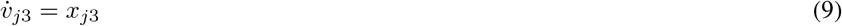

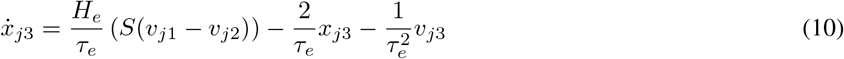

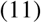

where *v*_1,2,3_ is the average postsynaptic membrane potential of the excitatory interneuron, inhibitory interneuron, and pyramidal cell populations, respectively. The input from other nodes in the whole-brain network

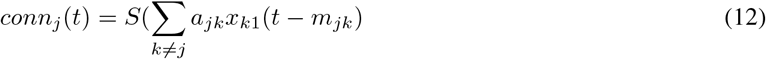

where *a*_*jk*_ is the *j*th row and the *k*th column in the connectivity matrix **A** (which in our is the rescaled connectivity Laplacian as described above). conn_*j*_ thus enters into the excitatory population only and collects excitatory population activity from other network nodes. Due to the finite velocity of long-range axonal conduction, these inputs appear after delays of around 5-50ms, which vary on a per-connection basis. This is specified by *m*_*jk*_, the *j, k*th entry of the delays matrix **M** = **T***/s*, which is a function of the inter-regional fibre tract length matrix **T** and the global axonal conduction velocity *s*. Especially important here, the TMS-induced depolarization of the resting membrane potential was modelled by an external perturbing voltage offset *p* applied to the excitatory interneuron population.

To establish which parcels in the model the TMS stimulation is injected into, and with what strength, the TMS-induced electric field was modelled with SimNIBS^63^ in the MNI152 standard-space. The normalized electric field or *E-field* distribution was thresholded at 83% of its maximal value, following recent estimates of the E-field thresholds above which tissue is activated by TMS^64^. This thresholded E-field map was then used to inject a weighted stimulus into the target regions in the model. Finally, channel-level EEG signals were computed in the model by first taking the difference *y*(*t*) = *v*_1_(*t*) − *v*_2_(*t*) between the excitatory and inhibitory interneuron activity at each cortical parcel^65^, and projected to the EEG channels space using a leadfield matrix.

### 4.5 Individual-subject Jansen-Rit connectome model parameter estimation from TMS-EEG data

We used a novel brain network model parameter optimization technique^24^ for fitting individual-subject TEP waveforms and identifying subject-level physiological parameters from empirical data. Notably, the model is implemented in PyTorch^40^, a software library that has in recent years been widely adopted by the machine learning community in both academic and commercial sectors. Moving to this framework from more conventional numerical simulation libraries involves some minor modifications to accommodate tensor data structures, but brings the substantial advantage of naturally accommodating gradient-based parameter optimization via automatic differentiation-based algorithms, for relatively complex sets of equations that do not admit of tractably computable Jacobians. This is one of a growing number of cases (e.g.^66;67^) where the natural parallel between our physiologically-based large-scale brain network models and deep recurrent neural networks used in machine learning is proving technically and conceptually fruitful. The general mathematical framework for this approach has been described by us in a recent technical paper^24^, where it was applied in the context of connectome-based neurophysiological modelling of resting-state fMRI data. In the present work we are extending this technique’s domain of application to fast-timescale evoked responses, but the overall approach in the two cases is the same with minor modifications. The algorithm proceeds by dividing a subject’s multi (in this case 64) -channel, 600ms long (−100ms to +500ms post-stimulus), trial-averaged TMS-EEG TEP waveform into short (40ms) non-overlapping windows, termed *batches*. Rolling through each batch in the time series sequentially, the JR model-simulated TEP **ŷ** was generated with the current set of parameter values, and its match to the empirical TEP **y** was calculated with the following mean-squared error (MSE) loss function

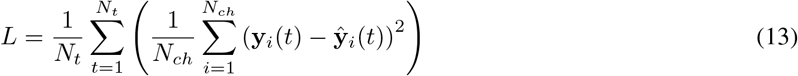

where *N*_*t*_ is the number of the timepoints and *N*_*ch*_ is the number of EEG channels. It is assumed that model parameters are Gaussian. Together with a complexity-penalizing regularization term on each model parameter *θ*,

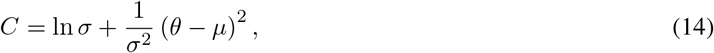

where the mean *µ* and standard variation *σ* of the model parameter *θ* are hyper-parameters to be fitted. The model parameters’ complexity defined in Eq (14) is included as a regularization term to avoid over-fitting and help achieve a robust model. The loss function *L* and the complexity term *C* are combined into a final objective function that is provided to PyTorch’s native ADAM algorithm^46^, which selects the candidate parameter set for the next batch with a stochastic gradient descent-based scheme that utilizes automatic-differentiation-based gradients (efficient computation of which is primary design objective of the (Py)Torch C++ backend). When the batch window reaches the end of the TEP time series, it returns to the start and repeats until convergence. For an overview of all parameters used in the model, please refer to Supplementary Figure S7. For a complete description of the parameter estimation algorithm, please see^24^.

### 4.6 Assessing similarity between simulated and empirical TEPs

To further assess goodness-of-fit of the simulated TEP waveforms arrived at after convergence of the ADAM algorithm, we conducted additional analyses in both EEG sensor and source space. At the channel level, Pearson correlation coefficients and corresponding p-values between empirical and model-generated TEP waveforms were computed for each subject. In order to control for type I error, this result was compared with a null distribution constructed from 1000 time-wise random permutations, with a significance threshold set at *p <* 0.05. As a complement to these TEP comparisons that emphasize matching of waveform shape and component timing, we also examined more holistic time series variability characteristics using the PCI^27^, which was extracted from the simulated and the empirical TMS-EEG data, and Pearson correlations between the two computed. Assessment of goodness-of-fit at the source level proceeded in a similar fashion: Individual subjects’ empirical and model-generated TMS-EEG timeseries were first computed for every source-space surface vertex, as described above. Pearson correlation coefficients and corresponding p-values, indicating empirical-simulated data similarities, were computed. Again, in order to control for type I error, time-wise permutation testing was done by comparison against 1000 surrogate, shuffled TEP differences, with a significance threshold set at *p <* 0.05. Finally, and unlike the channel-level data, network-level comparisons of simulated vs. empirical activity patterns were made by averaging current densities over surface vertices at each point in time within each of the 7 Freesurfer surface-projected canonical Yeo network maps^28^, and Pearson correlation coefficients and p-values between empirical and simulated network-level time series were again computed.

### 4.7 Dissecting the propagation dynamics of TMS-evoked responses

A key aim of the present study is to ascertain whether the TMS-evoked activity in a certain region at a certain time point is primarily attributable to a localized response to TMS at the primary stimulation site, or to re-entrant activity feeding back from other nodes in the connectome network. In order to explore this, activity of each network node at a given time point was extracted as the sum of the absolute value of the simulated pyramidal cell population activity within a narrow temporal window (0-300ms). Maximally activated nodes were defined as the top 1% of nodes exceeding two standard deviations above the mean over regions. This approach was used to identify, for each subject individually, the most important nodes at three key time points: 20ms, 50ms, 100ms after the TMS pulse, where we wanted to identify the contribution of re-entrant activity.

With these key brain regions identified for each time window of interest, simulations were re-run for each subject using their optimal parameters estimated from the original TEP fitting step - but this time with the selected nodes’ incoming and the outgoing connection weights set to zero for the duration of the window. These new ‘virtually lesioned’ TEP time series were again projected to the EEG channel space and back to the source level, where they were compared against the original model-generated TEP time series. Finally, as above, the model-generated dSPM values were extracted from the 7 canonical network surface maps for each individual and for each condition, and analyzed statistically using the Statistical Package for the Social Sciences (SPSS) Version 25 (IBM Corp). A 4×7 repeated measures ANOVA with within-subjects factors “TIME OF DAMAGE (4 levels: 20ms; 50ms; 100ms; no damage) and “NETWORK” (7 levels: VISN; SMN; DAN; ASN; LIMN; FPN; DMN) was run. Post-hoc paired t-tests were used to detect dSPM value changes for different networks and lesion times, testing on a per-network basis whether and at what times the virtual lesions impacted on network-level activations.

### 4.8 Evaluating how the anatomical connectome affects TMS-evoked EEG dynamics

Complementing the analyses probing the importance of incoming and outgoing activity of the most-active regions at key TEP timepoints, we also performed the same time-windowed virtual lesion analyses for regions based on their role in the brain network’s graph structure. This provided additional insight into the importance of the anatomical connectome in shaping the propagation of the TMS-evoked signal. In order to do this, the out-degree *O*_*i*_ of every node *i* in the original (prior) tractography-derived connection weights matrix **A** was calculated as the number of outgoing edges

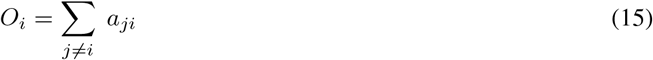

where *a*_*ij*_ is the element of the ith row and jth column of **A**, and the sum is over all nodes in the network. In the following we then focused on the top 1% of nodes according to out-degree. Simulation of these virtual lesions proceeded exactly as above but with the incoming and the outgoing connections of the selected nodes set to zero at different time point depending on the conditions (e.g. 20ms, 50ms or 100ms after the external input). In graph theory and network science, investigation of network properties by lesioning the most important nodes in this way is known as a ‘targeted attack’^68^. As a corresponding ‘random attack’ control condition^69^, 1000 simulations were also run where the nodes selected for removal of their incoming and outgoing collections were randomly chosen. For both random and targeted attack simulation conditions, the anatomical connectome was damaged at the same set of key time points - 20ms, 50ms, and 100ms after the TMS pulse. To assess these comparisons statistically, we first examined PCI extracted from the simulated TMS-EEG time series. This was done using a 2×3 repeated measures ANOVA with within-subjects factors “ATTACK TYPE” (2 levels: targeted attack; random attack) and “TIME OF DAMAGE” (3 levels: 20ms; 50ms; 100ms). Post-hoc paired t-tests were used to examine dSPM values changes for individual types and times of damage. Finally, the simulated TMS-EEG time series were projected into source space, and dSPM values were extracted from the 7 Yeo network maps for each conditions. A 2×3×7 repeated measures ANOVA with within-subjects factors “ATTACK TYPE” (2 levels: targeted attack; random attack) and “TIME OF DAMAGE” (3 levels: 20ms; 50ms; 100ms) and “NETWORK” (7 levels: VIS; SMN; DAN; ASN; LIM; FPN; DMN) was then used to test for key effects of interest. Subsequently, post-hoc paired t-tests were used to detect dSPM value changes for different networks and different types and times of damage.

### 4.9 Identifying clusters of different TMS-evoked responses

We were aimed at predicting the spatiotemporal propagation of the TMS-evoked signal using the optimized physiological parameters of the model. Firstly, singular value decompositions (SVDs) were run on the grand mean of both the empirical and the model-generated TMS-EEG data, in order to identify prototypical TMS-evoked responses. Following this, the group-level SVD spatial eigenmodes were identified within each subject’s time series corresponding to the time point with the highest cosine similarity between the individual’s TEP and the prototypical response. Latencies and amplitudes of the SVD left singular vector time series peaks were extracted for every subject and related with the individuals’ JR model parameters, with Pearson correlation coefficients and corresponding p-values were computed accordingly.

## Code and Data Availability

Full code for reproduction of the data analysis and model fitting described in this paper is freely available online at github.com/GriffithsLab/PyTepFit. As noted above, TMS-EEG data were taken from an open dataset (figshare.com/articles/dataset/TEPs-_SEPs/7440713). Structural MRI data used in the study are available from the original Human Connectome Project dataset (`humanconnectome.org`).

## Notes

https://github.com/GriffithsLab/PyTepFit

### Competing Interest Statement

The authors have declared no competing interest.

