## Supplementary Information for "TMS-Evoked Responses Are Driven by Recurrent Large-Scale Network Dynamics"

May 31, 2022

### Contents

**1 Supplementary Materials and Methods** **3**

1.1 Modelling of the TMS-induced electric field . . . . . . . . . . . . . . . . . . . . . . . . . . . . . . 3

**2 Supplementary Results** **4**

2.1 Statistical analyses of TMS-evoked activity propagation and maximum activation-based virtual lesions 4

2.2 Statistical analyses of TMS-evoked activity propagation and maximum network topology-based virtual lesions . . . . . . . . . . . . . . . . . . . . . . . . . . . . . . . . . . . . . . . . . . . . . . 4

**3 Supplementary Figures** **5**

### 1 Supplementary Materials and Methods

#### 1.1 Modelling of the TMS-induced electric field

In order to identify the brain regions engaged by M1-targeted TMS, and therefore which nodes in our simulation to inject an external input, and with what magnitude, the TMS-induced electric field was modelled with SimNIBS[1] (`simnibs.github.io/simnibs`). A tetrahedral head model (mesh file) was created, consisting of five tissue types: white matter (WM), grey matter (GM), cerebro-spinal fluid (CSF), skull, and scalp. The assigned conductivity values were fixed: 0.126 S/m (WM), 0.275 S/m (GM), 1.654 S/m (CSF), 0.01 S/m (skull), 0.465 S/m (scalp). The distance between the coil and the cortex was set to 10 mm, as measured in our MRI images, and the coil handle was oriented following the M1 coordinates following the methods and materials of the original paper of Bibiani et al.[2]. The rate of change of the coil current (dI/dt) was calculated assuming a quasi-static regime[3], according to the following equation:

$$\mathbf{E} = \frac{\vartheta \mathbf{A}}{\vartheta t} - \Delta\varphi$$

where $\mathbf{E}$ is the electric field vector, $\varphi$ denotes the electric potential, and $\mathbf{A}$ is the magnetic vector potential of the TMS coil, which depends on the coil's shape, position, and the current flow in the coil wires.

### 2 Supplementary Results

#### 2.1 Statistical analyses of TMS-evoked activity propagation and maximum activation-based virtual lesions

Continuing the results reported in main text Section 2.2 from ANOVA on network ROI-averaged source activity (dSPM) values, we conducted the following post-hoc t-tests to compare simulated network activation by TMS for different networks and virtual lesion timings: Taking into account the factor TIME OF DAMAGE, post-hoc comparisons showed significant differences in dSPM values for: 20ms>50ms: mean difference=-0.52, p<0.0001; 20ms>100ms: mean difference=-2.16, p<0.0001; 20ms>no damage: mean difference=1.53, p<0.0001; 50ms>100ms: mean difference=-1.63, p<0.0001; 50ms>no damage: mean difference=1.01, p<0.0001; 100ms>no damage: mean difference=0.62, p<0.0001). Conversely, considering the factor NETWORK, post-hoc comparisons showed significant differences in dSPM values for: VIS>SMN: mean difference=-1.91, p<0.0001; VIS>DAN: mean difference=-0.90, p<0.0001; VIS>ASN: mean difference=-1.93, p<0.0001; VIS>LIM: mean difference=-0.21, p=0.002; VIS>FPN: mean difference=-0.79, p<0.0001; VIS>DMN: mean difference=-0.72, p<0.0001; SMN>DAN: mean difference=1.01, p<0.0001; SMN>ASN: mean difference=1.72, p<0.0001; SMN>LIM: mean difference=2.12, p<0.0001; SMN>FPN: mean difference=1.11, p<0.0001; SMN>DMN: mean difference=1.19, p<0.0001; DAN>ASN: mean difference=0.70, p<0.0001; DAN>LIM: mean difference=1.11, p<0.0001; DAN>DMN: mean difference=0.18, p=0.006; ASN>LIM: mean difference=-0.40, p<0.0001; AS>FPN: mean difference=0.60, p<0.0001; ASN>DMN: mean difference=-0.52, p<0.0001; LIM>FPN: mean difference=-1.01, p<0.0001; LIM>DMN: mean difference=-0.93, p<0.0001; FPN>DMN: mean difference=-0.07, p=0.002). Conversely no significant differences were found for DAN>FPN: mean difference=0.11, p=0.1. These post-hoc comparisons indicate that first, early damages of the structural connectome (e.g. 20ms and 50ms) compromise the propagation of the TMS-evoked signal compare to the other conditions (e.g. 100ms and no damage). Second, the network where this result is more pronounced is the stimulated SMN.

#### 2.2 Statistical analyses of TMS-evoked activity propagation and maximum network topology-based virtual lesions

Again, continuing the results reported in main text Section 2.3 from ANOVA on network ROI-averaged source activity (dSPM) values, the following post-hoc t-tests were conducted to compare simulated network activation by TMS in the presence of virtual lesions targeted to highly connected regions vs. virtual lesions selected randomly. Considering the factor TIME OF DAMAGE, post-hoc comparisons showed significant differences in dSPM values for: 20ms>50ms: mean difference=-0.63, p=0.001; 20ms>100ms: mean difference=-0.17, p<0.0001; 50ms>100ms: mean difference=-0.11, p<0.0001). Conversely, taking into account the factor ATTACK TYPE, post-hoc comparisons revealed significant differences in dSPM values comparing targeted>random: mean difference=-0.32, p<0.0001. Finally, considering the factor NETWORK, significant differences in post-hoc comparisons were found for: VIS>SMN: mean difference=-0.28, p<0.0001; VIS>DAN: mean difference=-0.08, p<0.0001; VIS>ASN: mean difference=-1.54, p<0.0001; VIS>FPN: mean difference=-0.17, p<0.0001; VIS>DMN: mean difference=-0.26, p<0.0001; SMN>DAN: mean difference=0.21, p<0.0001; SMN>ASN: mean difference=0.13, p<0.0001; SMN>LIM: mean difference=0.31, p<0.0001; SMN>FPN: mean difference=0.11, p<0.0001; DAN>ASN: mean difference=-0.07, p=0.009; DAN>LIM: mean difference=-0.09, p=0.002; DAN>FPN: mean difference=-0.09, p<0.0001; DAN >DMN: mean difference=-0.18, p<0.0001; ASN>LIM: mean difference=0.16, p=0.009; ASN>DMN: mean difference=-0.11, p<0.0001; LIM>FPN: mean difference=-0.18, p<0.0001; LIM>DMN: mean difference=-0.28, p<0.0001; FPN>DMN: mean difference=-0.09, p=0.01). On the other hand, no significant post-hoc comparisons were found for: VIS>LIM: mean difference=0.01, p=0.69; SMN>DMN: mean difference=0.02, p=0.31; ASN>FPN: mean difference=-0.02, p=0.28.The main result of these post-hoc comparisons is that both space and time of the lesions are important for significantly affecting the propagation of the TMS-evoked activity.

### 3 Supplementary Figures

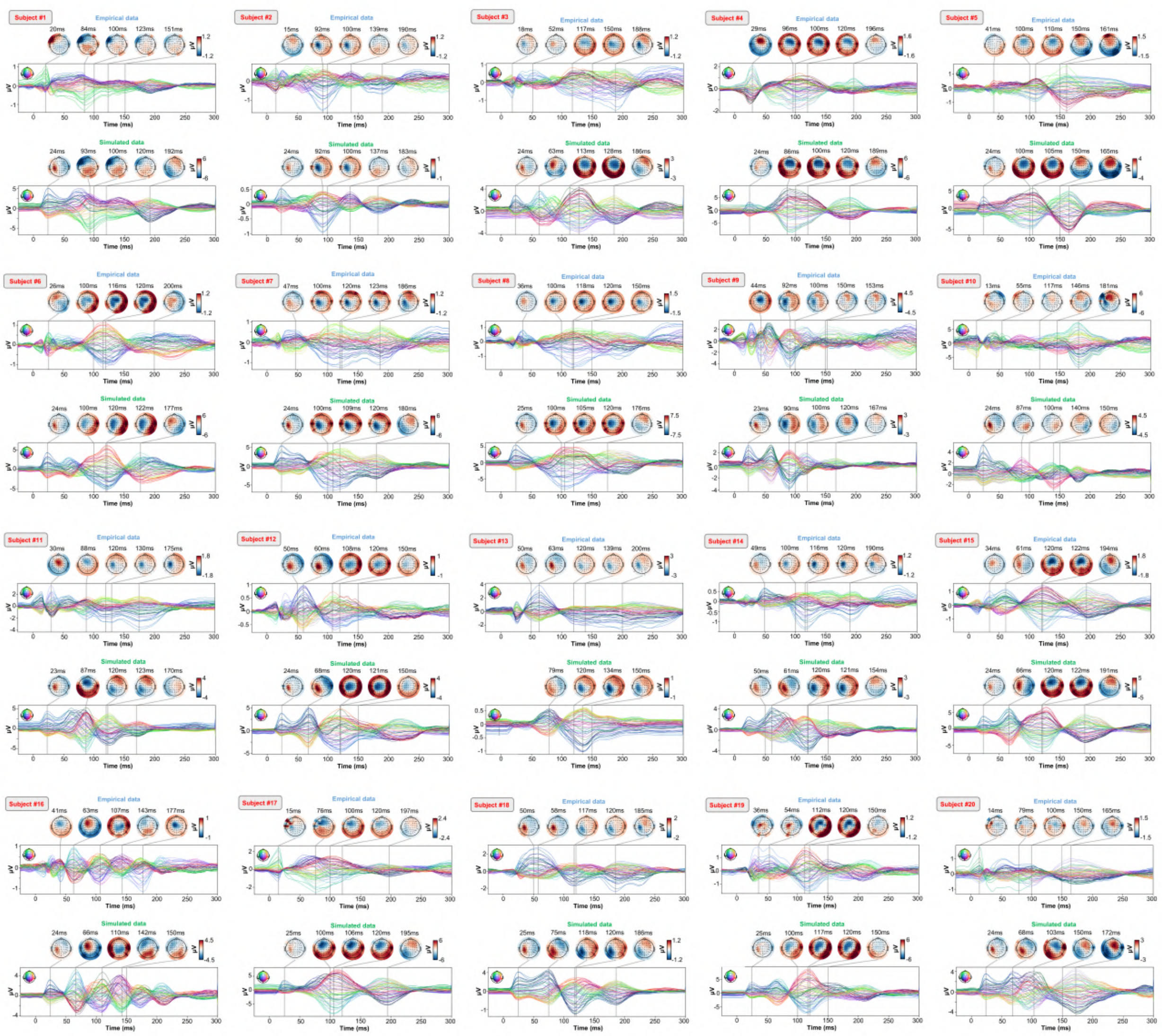

**Figure S1 |** ***Optimized TEP models for all subjects.*** For every pair of rows, empirical (upper) and simulated (lower) TMS-EEG responses are shown for every study subject, extending main text Figure 2 where a selected subset of subjects' data are shown. These data reiterate and reinforce the demonstrations in Figure 2 that the model-generated EEG activity time series achieve robust recovery of individual subjects' empirical TEP propagation patterns.

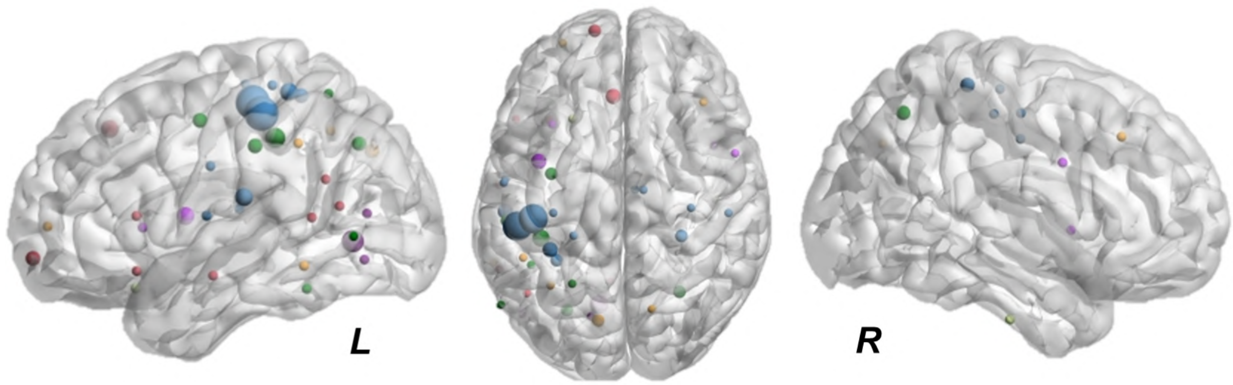


**Figure S2 |** ***Representation of the nodes removed during the analysis.*** Sizes indicate for how many subjects a node was removed. A clear pattern is evident, where the more important nodes are the ones more affected by the stimulation (e.g. motor, parietal and frontal cortex).

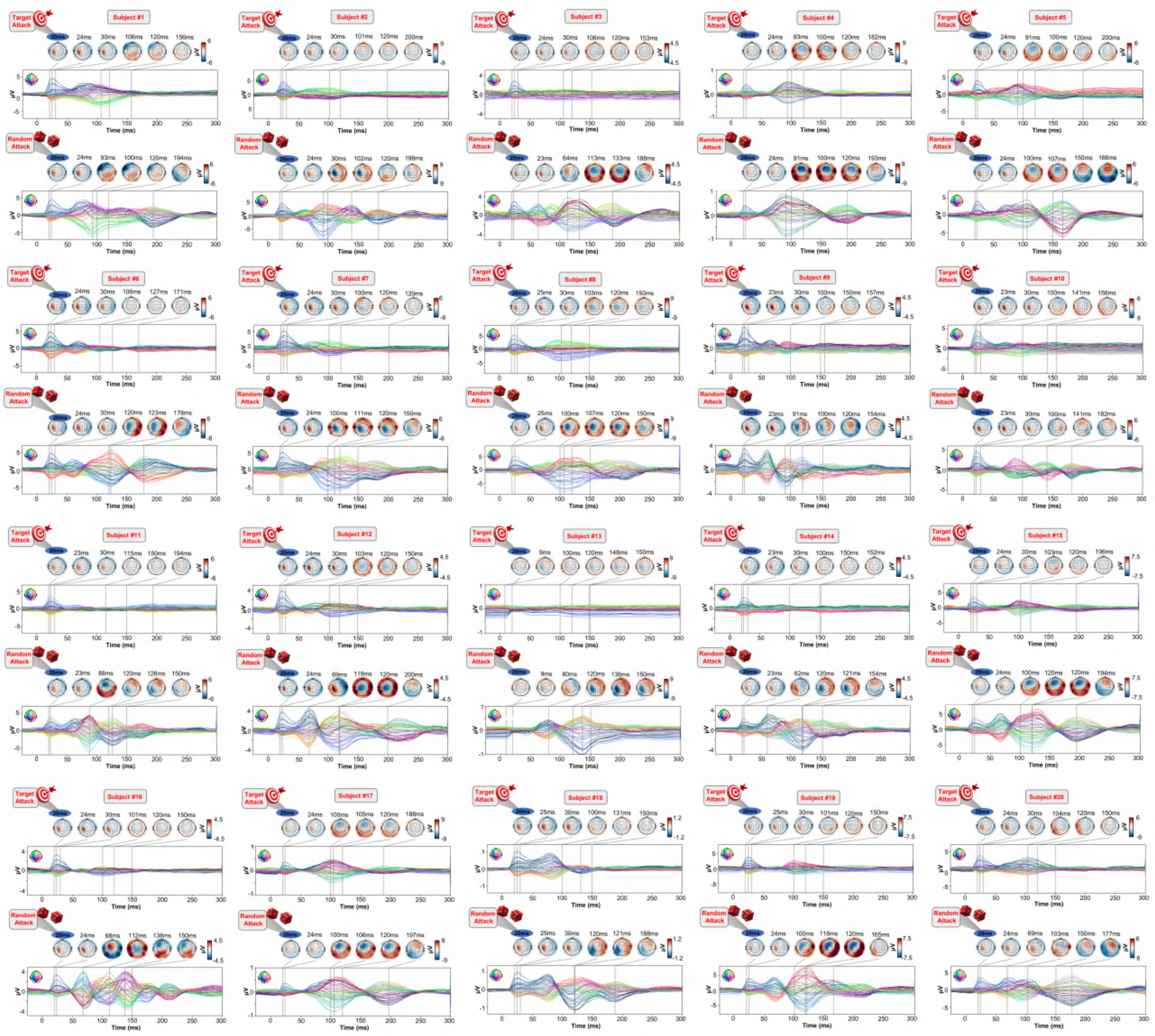

**Figure S3 |** ***TEP models for all subjects with targeted and random connectome-based virtual lesions at 20ms.*** Extending main text Figure 4 to all study subjects: For every pair of rows, simulated TMS-EEG responses confirm that targeted attack (upper) significantly compromised the propagation of the TMS-evoked activity, whereas random attack (lower) had little effect, resulting in TEPs similar to the original no-damage models in Figure S1.

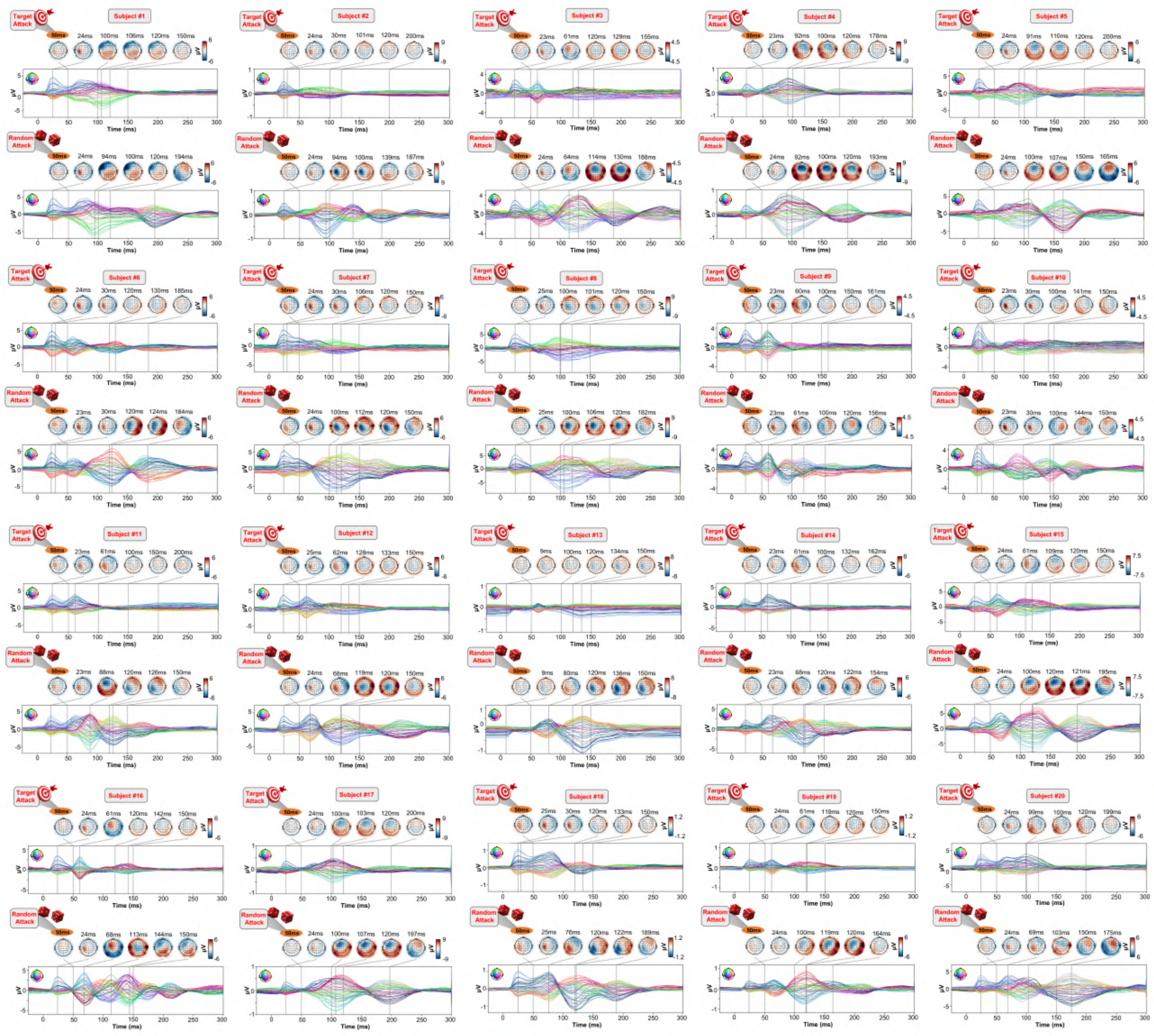

**Figure S4 | *TEP models for all subjects with targeted and random connectome-based virtual lesions at 50ms.*** Follows same structure to Figure S2.

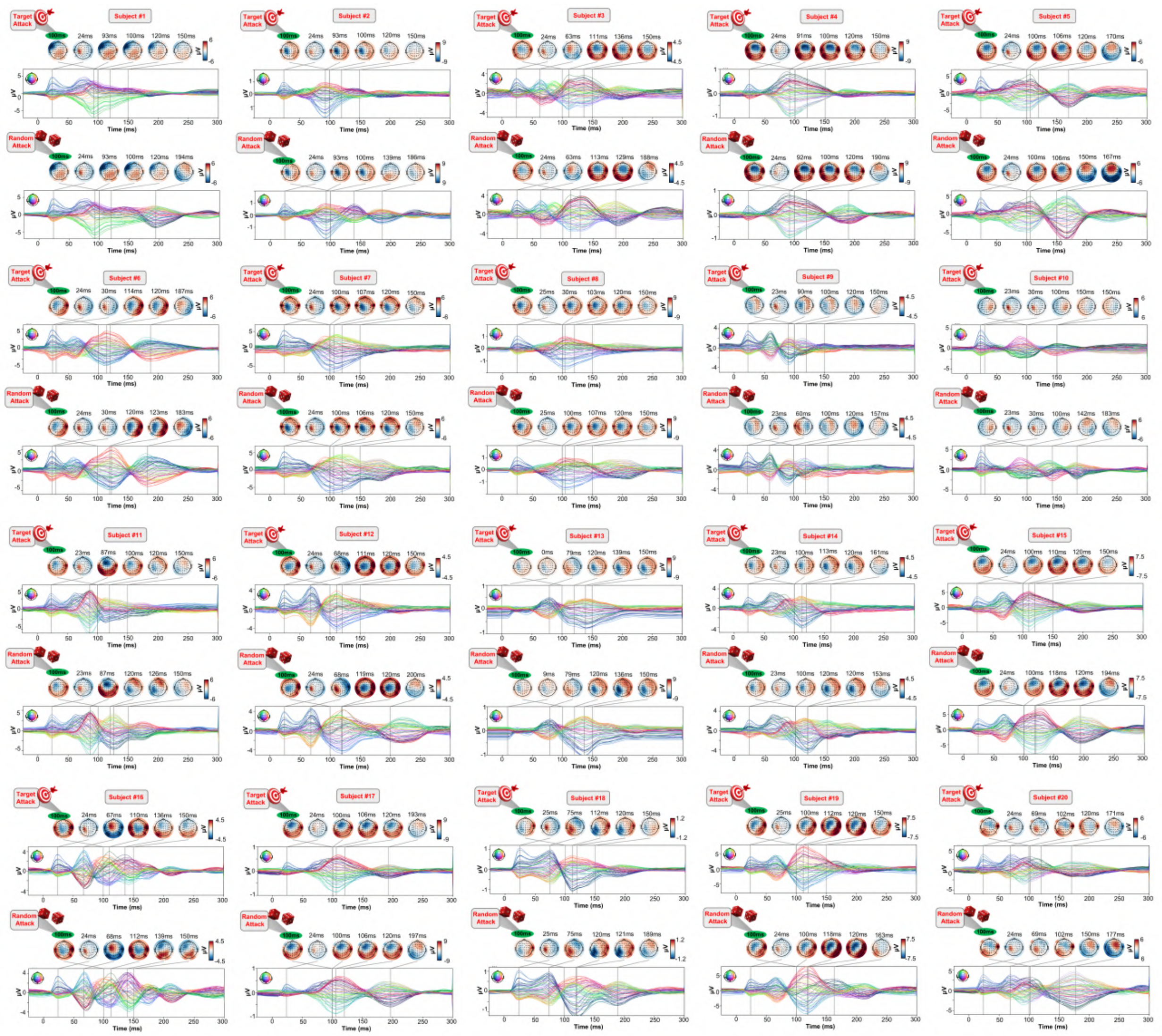

**Figure S5 |** ***TEP models for all subjects with targeted and random connectome-based virtual lesions at 100ms.*** Follows same structure to Figure S2.

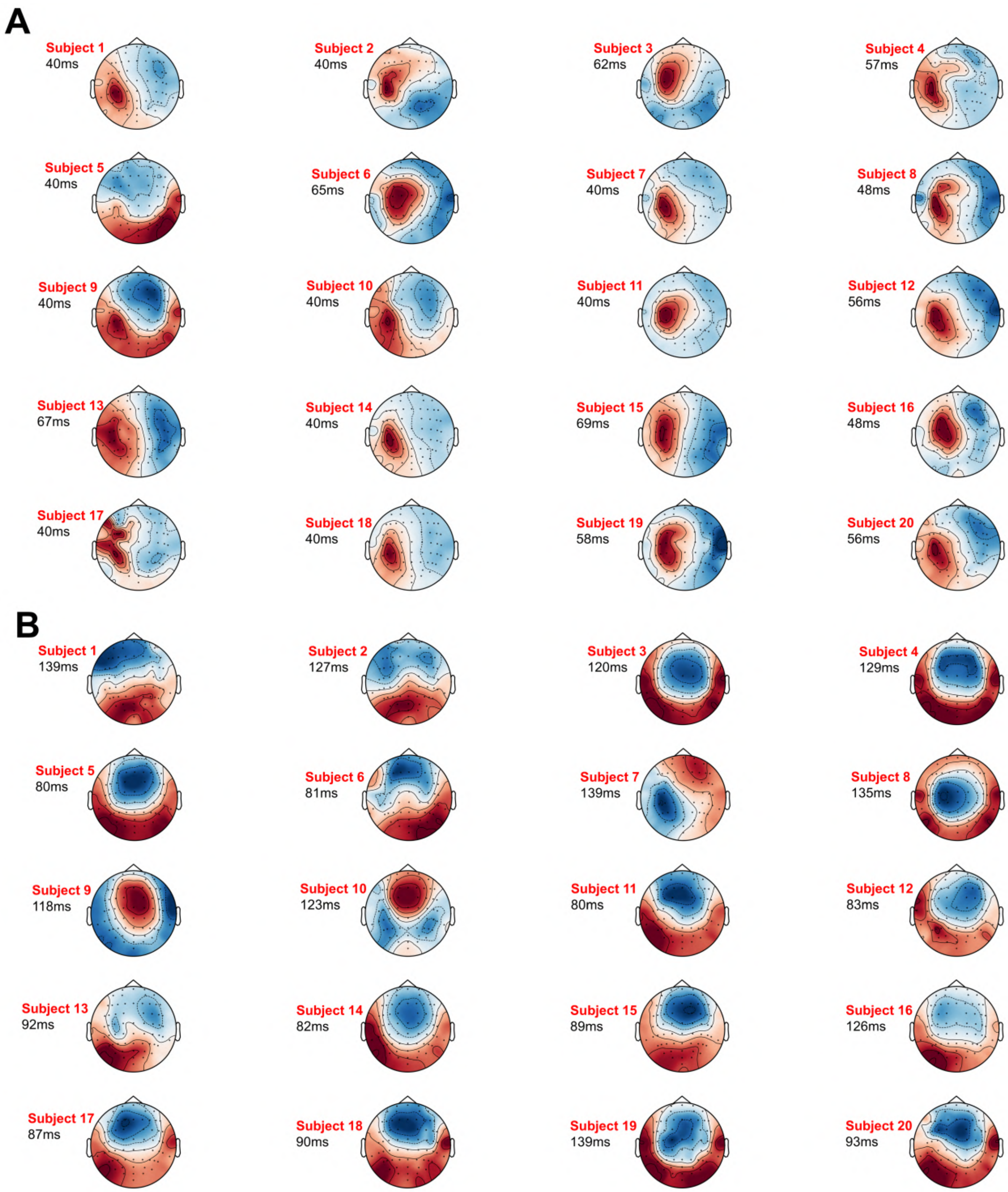


**Figure S6 | *Timing and topographies of the prototypical TEP response pattern in each subject.*** These figures extend the single-subject examples from TEP channel data SVD decompositions in Figure 6. **A)** First right singular vectors from TEP SVDs for all subjects, with corresponding time location indicating the time point of maximum expression for the corresponding left singular vector (temporal eigenmode). **B)** Second right singular vectors and corresponding time points of maximum expression.

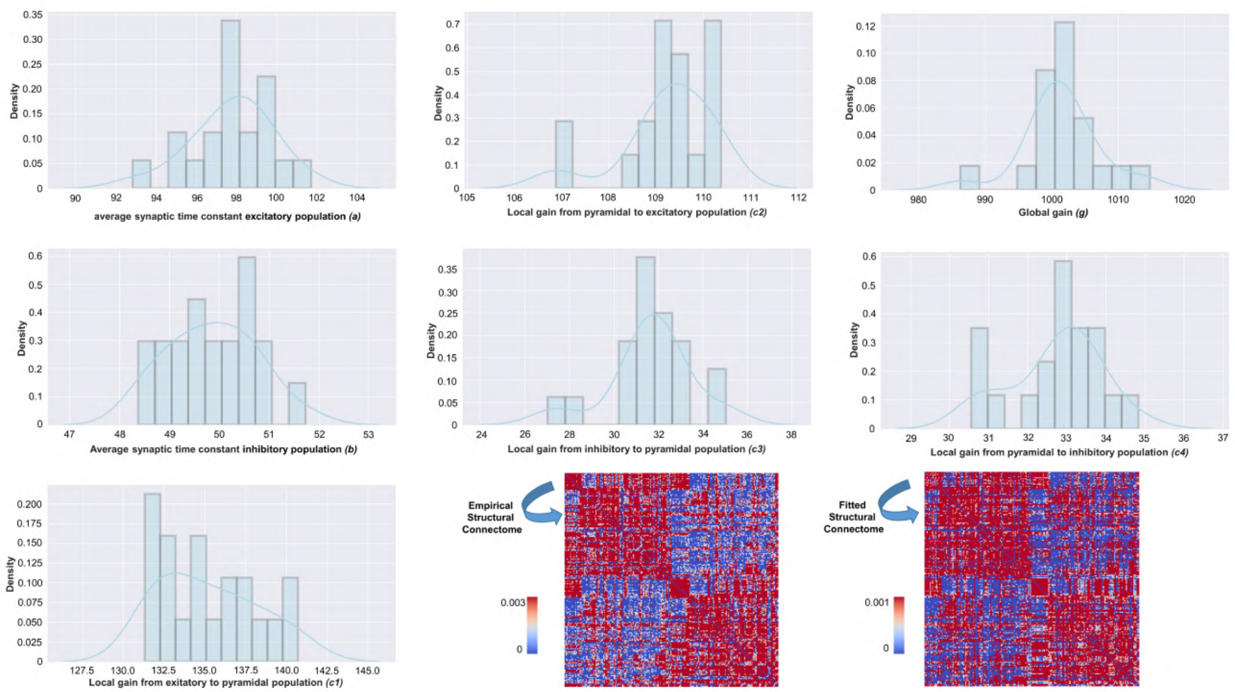


**Figure S7 |** ***Distributions of physiological parameter estimates over subjects.*** Histograms and kernel density estimates of the estimated values for the Jansen-Rit model physiological parameters over all subjects. Also shown are prior and posterior parameter values for anatomical connectome weights for a single example subject (bottom right). Parameter estimation was performed using our novel automatic differentiation and gradient-based approach inspired by current techniques in deep learning[4].

[4] Deep learning-based parameter estimation for neurophysiological models of neuroimaging data. .
